# Comparing and Validating Automated Tools for Individualized Electric Field Simulations in the Human Head

**DOI:** 10.1101/611962

**Authors:** Oula Puonti, Guilherme B. Saturnino, Kristoffer H. Madsen, Axel Thielscher

**Affiliations:** Danish Research Centre for Magnetic Resonance, Centre for Functional and Diagnostic Imaging and Research, Copenhagen University Hospital Hvidovre, Hvidovre, Denmark; Department of Health Technology, Technical University of Denmark, Kgs. Lyngby, Denmark; Department of Applied Mathematics and Computer Science, Technical University of Denmark, Kgs. Lyngby, Denmark

**Keywords:** transcranial brain stimulation, TDCS, TACS, volume conductor model, errors-in-variables regression, Bayesian regression

## Abstract

Comparing electric field simulations from individualized head models against in-vivo recordings is important for direct validation of computational field modeling for transcranial brain stimulation and brain mapping techniques such as electro- and magnetoencephalography. This also helps to improve simulation accuracy by pinning down the factors having the largest influence on the simulations. Here we compare field simulations from four different automated pipelines, against intracranial voltage recordings in an existing dataset of 14 epilepsy patients. We show that ignoring uncertainty in the simulations leads to a strong bias in the estimated linear relationship between simulated and measured fields. In addition, even though the simulations between the pipelines differ notably, this is not reflected in the correlation with the measurements. We discuss potential reasons for this apparent mismatch and propose a new Bayesian regression analysis of the data that yields unbiased estimates enabling robust conclusions to be reached.

## INTRODUCTION

Modeling the current flow distribution in the brain is in the core of many neuroimaging and functional brain mapping techniques. The currents-of-interest can be either externally induced by transcranial brain stimulation (TBS) methods, such as transcranial electrical stimulation (TES) or transcranial magnetic stimulation (TMS) or they can result from neuronal activity in which case they can be measured using potential differences on the scalp (EEG) or by recording the produced magnetic fields (MEG). In TBS, the individual anatomy has a large, and often counter-intuitive, impact in shaping the current flow inside the cranium (Bungert et al., 2017; Datta et al., 2009; Laakso et al., 2015; Miranda et al., 2013; Opitz et al., 2015). Similarly, the signal measured in EEG and MEG is dependent on the complex geometry of the head (Cho et al., 2015; Dannhauer et al., 2011; Stenroos et al., 2014). These findings have prompted a shift away from simplified anatomical models (Ravazzani et al., 1996) towards individualized head models based on structural magnetic resonance (MR) scans. Individualized modeling holds great promise particularly for transcranial electric stimulation (TES), where results from stimulation studies (López-Alonso et al., 2014; Wiethoff et al., 2014) show large inter-subject variability in responses to stimulation. Part of this variability is likely explained by dosing differences due to anatomical variation. Individualized modeling enables dose control and can be used to systematically improve spatial targeting by automated tailoring of the electrode positions and injected currents, ensuring that the highest field strengths are contained to the region-of-interest (Dmochowski et al., 2011). This opens the door for personalized treatment approaches in a variety of brain disorders ranging from major depressive disorder (Csifcsák et al., 2018) to motor rehabilitation after stroke (Datta et al., 2011; Minjoli et al., 2017).

Although practically relevant results were obtained from simulation studies that support the usefulness of individualized head models, one of the key challenges is the direct in-vivo validation of the electric field simulations in the human brain. The modeling process includes uncertainties, mainly related to the segmentation of the anatomy (Nielsen et al., 2018) and spatial tissue conductivities (Saturnino et al., 2019), which propagate onto the reconstructed fields. Optimally, one would use in-vivo field measurements in the brain to gauge the accuracy of the simulations. In practice this is, however, difficult: In-vivo measurements of the electric fields are experimentally very demanding and susceptible to measurement errors, creating unwanted uncertainty in the data which is supposed to be used as ground truth for validating the simulations. To-date, we are aware of two studies where head model validation using direct in-vivo intra-cranial measurements of the electric fields in humans is attempted. The first one by (Huang et al., 2017) reports TES-induced voltage measurements on ten epilepsy patients with intra-cranial electrodes. The authors use the voltage measurements for assessing the correlations between the simulated and measured voltage differences and for calibrating the tissue conductivities of the individual head models. In similar vein, (Opitz et al., 2018) compare simulated and measured fields in two epilepsy patients reporting slightly lower correlations compared to those in (Huang et al., 2017).

The difference is probably explained by differences in the experimental procedures, as the recording and TES electrodes were quite close to each other and to nearby skull defects in the study of (Opitz et al., 2018), suggesting that discrepancies between real and modeled positions and anatomy might have had stronger effects on the field comparisons than in (Huang et al., 2017). (Huang et al., 2017) found that models based on “standard” literature values for the ohmic conductivities systematically overestimated the recorded voltage differences. On the other hand, (Opitz et al., 2018) found underestimated fields in one of the studied patients, while the calculated e-fields were too high in the second patient. Recently, (Göksu et al., 2018) demonstrated a novel non-invasive approach to reconstruct TES induced current densities in the brain from magnetic resonance images of the current-generated magnetic fields (magnetic resonance current density imaging, MRCDI). They present initial results on five subjects showing good agreement between simulated and measured current densities, with a moderate but systematic underestimation of the current densities by the models based on “standard” ohmic conductivities. Non-invasive measurements would be the preferred approach, not only due to ethical aspects relating to human studies, but also because invasive measurements change the volume conduction properties of the head, thus introducing additional modeling complexities. MRCDI is a promising step to the correct direction but needs further development before it can be applied for head model validation.

In this article, we compare four different automated methods for end-to-end electric field simulations starting from a structural MR scan, followed by segmentation of the anatomy and generation of a finite element (FEM) mesh, and finally calculating the electric field distribution in the brain for a given stimulation protocol. We show that even though the simulated electric field distributions can differ notably between the methods, the correlations with the measured voltage differences can be very similar. We further show the importance of considering uncertainties in both the recorded *and* simulated potential differences in the analysis. The results highlight the difficulty of validating the simulations, even when direct measurements are available, and point to a need for a more careful analysis of the available data and for adopting a strategic approach to future measurement studies in order to reach conclusive validations. The analysis is based on the freely available data set from (Huang et al., 2017) and the results should be contrasted to those presented in (Huang et al., 2018), where significant differences between the methods in predicting the measurements were found.

## MATERIAL AND METHODS

The data set consists of 14 epilepsy patients with intracranial EEG electrodes planted for surgical evaluation (Huang et al., 2017). For each subject there exists a T1-weighted MR scan, a manually corrected segmentation of the main tissue classes (white matter - WM, gray matter - GM, cerebro-spinal fluid - CSF, skull and scalp) along with annotations of the stimulation electrodes, subgaleal electrodes, intracranial electrode strip, and the surgical drain. The locations, in MNI and voxel coordinates, and measured voltages from the intracranial electrodes are provided as a text file. The data set is freely available for download after registration at http://dx.doi.org/10.6080/K0XW4GQ1.

We compare two different software tools for generating individualized head models and simulating the electric fields induced by TES: *SimNIBS 2.1* (Saturnino et al., 2018) and *ROAST v2.7* (Huang et al., 2018). *SimNIBS 2.1* offers three alternative approaches for generating the anatomical head models, which we consider individually, giving in total four methods to compare. For completeness, we will next briefly describe each of the approaches.

### Head model generation

- The default pipeline for head model generation in *SimNIBS 2.1*, called *headreco* (Nielsen et al., 2018), uses the segmentation routine from *SPM12* (http://www.fil.ion.ucl.ac.uk/spm/software/spm12/) (Ashburner and Friston, 2005), combined with an extended anatomical atlas (Huang et al., 2013), to generate a tissue segmentation from a set of possibly multi-contrast MR scans. After the initial segmentation, the tissue masks are cleaned using simple morphological operations to reduce noise and ensure that the tissues are contained within each other. Next, surfaces, represented as triangular elements, are extracted from the voxel segmentations. As a last step, the FEM mesh is generated by filling in the space between the surfaces with tetrahedra. Note, that due to the chosen meshing approach, which first generates surfaces, the tissue classes need to be nested which is ensured by the clean-up step after the initial segmentation. For further details we refer the reader to (Nielsen et al., 2018).
- The *headreco* pipeline supports detailed cortical surface reconstructions, which are generated using the computational anatomy toolbox (*CAT12*, http://www.neuro.uni-jena.de/cat/) implemented in *SPM12*. The cortical surfaces from *CAT12* replace the ones generated from the voxel segmentations in the standard *headreco* pipeline, while other parts of the pipeline remain the same. We denote this approach *headreco+CAT*.
- The predecessor of *headreco* in *SimNIBS*, called *mri2mesh* (Windhoff et al., 2013), combines the cortical surfaces and subcortical segmentation generated by *FreeSurfer* (https://surfer.nmr.mgh.harvard.edu/) (Fischl et al., 2002), with extra-cerebral tissue segmentations from FSL’s brain extraction tool (https://fsl.fmrib.ox.ac.uk/fsl/fslwiki) (Pechaud et al., 2006). Similar to *headreco* with *CAT12*, the cortical surfaces from *FreeSurfer* are combined with surfaces created from voxel segmentations in the other tissues, and the final FEM mesh is obtained by filling in tetrahedra.
- Similar to the standard version of *headreco*, the *ROAST v2.7* toolbox (http://www.parralab.org/roast/) (Huang et al., 2018) generates the tissue segmentations using *SPM12*, with the same extended anatomical atlas, and applies morphological operations to clean the segmentations. The main difference between the methods is in the post-processing and FEM meshing approach: whereas *headreco* first creates surfaces, roast generates a tetrahedral volume mesh directly from the voxel segmentation using CGAL (Fabri and Teillaud, 2011) called through the *iso2mesh* (http://iso2mesh.sourceforge.net/cgi-bin/index.cgi) *(Fang and Boas, 2009) toolbox. The restriction of nested tissue classes is thus relaxed, and anatomical details can potentially be better captured if the initial volume segmentation is accurate. However, the reconstructions of the tissue boundaries can be less accurate as the volume segmentation is meshed directly.*

### Simulating the electric fields

*The electric field calculations are performed in SimNIBS 2.1* for the head models generated with its pipelines (*headreco, headreco+CAT* and *mri2mesh*) and in *ROAST v2.7* for the head models generated with *ROAST*. In general, the electrodes are placed medially over the frontal and occipital poles (Huang et al., 2017). The electrodes are modeled as a 2×2cm square (Huang et al., 2017), and the thickness of the rubber and gel layers is set to 3mm for both, based on an estimation from the manually corrected segmentation volumes. The current flowing through the electrodes is set to 1mA, and the polarity adjusted to fit the recordings from (Huang et al., 2017) so that the direction of current flow is consistent with the measured data. Tissue and electrode conductivities were set to the values reported in (Huang et al., 2017).

Both *SimNIBS 2.1* and *ROAST v2.7* use the *GetDP* (Geuzaine, 2007) software to calculate electric potentials using the FEM method with first order tetrahedral elements. However, the post-processing of the simulations differs: *ROAST* uses *GetDP* to calculate the electric fields in each mesh node, while *SimNIBS* has native post-processing functions calculating the electric field for each mesh tetrahedra. The post-processing in *SimNIBS* is more consistent with the mathematical formulation of the Finite Element Method, where gradients are defined element-wise instead of node-wise (Zienkiewicz, O.C., Taylor, R.L, Zhu, 2013), and yields more physically plausible results, as the electric field values are discontinuous across tissue interfaces (Geselowitz, 1967). When interpolating or gridding results, *SimNIBS* uses the original mesh grid, keeping geometric consistency, while *ROAST* uses the *TriScatteredInterp* function in MATLAB (MathWorks, 2019), which does not preserve the original mesh, and instead creates a new Delaunay triangulation where the gridding is performed. Thus, the electric field values interpolated in *ROAST* do not observe tissue boundaries, as they do in *SimNIBS*.

## EXPERIMENTS

We performed two sets of experiments: the first one to quantify the differences in anatomical segmentation accuracy along with the differences in the simulated electric field distributions between the methods, and the second one to relate the electric field simulations to the measured potential differences in the intra-cranial electrodes. All the pipelines were run with default settings with the following exceptions:

- Both *headreco* and *headreco+CAT* were run with the *–d no-conform* option to avoid resampling of the input scans.
- For P04 in *headreco+CAT*, we set the vertex density (-v option) to 1.5 Nodes/mm^2^
- For P014 in *mri2mesh*, we set the number of vertices (--numvertices option) to 120000.
- For P010 the MR scan was resampled to 1mm^3^isotropic as *ROAST v2.7* does not account for anisotropic scans resulting in erroneous electric field estimates by effectively changing the electric conductivities along the axis where the anisotropy occurs. The developers of *ROAST* have fixed this bug in the newer version (2.7.1) after we pointed it out.
- For P06, we inverted the “x” component of the electric field calculated with *ROAST* to account for the fact that *ROAST* does not correct for the “x” axis flipping indicated by the header in the NifTi image.

The changes to the vertex densities in P04 and P014 were made as, after running the head model pipeline, we found that the head meshes were missing volumes (WM in P04 and CSF in P014). In both cases, increasing mesh density made surface decoupling more accurate and thus solved the problems in meshing the surfaces.

### Experiment 1

To assess the anatomical segmentation accuracy of the four methods, we compared the automatically generated tissue segmentations to the manually corrected ones using the Dice overlap score. The Dice score is defined as:

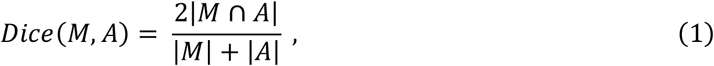

where *M* and *A* denote the manual and automated segmentation masks of a given tissue. We note that the manually corrected segmentations were created by first running the *ROAST* segmentation tool on the T1-weighted scans, then automatically correcting the output using a custom script, and finally correcting the remaining errors by hand (Huang et al., 2017). Some of the manually corrected segmentations still have inaccuracies (see Figure 2), and in general could be biased towards the automated *ROAST* segmentation. The Dice scores still serve as an indication of the general segmentation differences between the four methods. For the comparison, non-brain structures, such as the intra-cranial electrode strip and the surgical drain, were excluded from the manually corrected segmentations.

The differences between the simulated electric fields given by the methods were measured calculating the relative difference in the fields in GM for each pair of methods. The relative field difference is defined as:

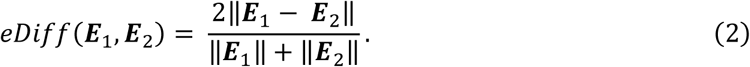

The differences were calculated using the electric fields transformed back from the FEM mesh into the original image grid using either *SimNIBS* or *ROAST*. In order to ensure an unbiased comparison, we calculated the relative field differences only where the gray matter mask in the two models under comparison overlap. This way we are sure that we compare the simulated electric fields within the same tissue, and avoid overestimation of the electric field differences due to differences in the segmentations. However, with this approach we are likely to underestimate the differences between the methods.

### Experiment 2

To relate the intracranial voltage recordings to the field simulations, we calculated the electric field along each recording electrode by taking the potential difference between neighboring contacts, divided by their distance. Similarly, we sampled the simulated voltages at the contact locations to calculate a corresponding electric field estimate for each of the automated methods. In *SimNIBS*, the sampling was performed using the electric potentials defined in the mesh nodes, while in *ROAST* it was done using the voltages transformed back to the original image grid.

Next, we fitted a standard linear model for each subject and method:

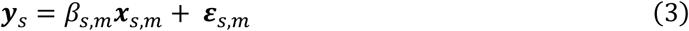

where ***x***_*s,m*_is a vector of the simulated potential differences for subject *s* and method *m*, ***y***_*s*_is a vector of the recorded potential differences in subject *s*, and the noise *ε*_*s,m*_for each subject and method is assumed to be normal i.i.d. For each subject and segmentation, we report the slope *β*_*s,m*_, the coefficient of determination (r^2^) and correlation (*ρ*). The same analysis was performed by (Huang et al., 2017) to compare simulations to intracranial electrode recordings.

We further extend this analysis by a hierarchical Bayesian regression analysis, which allows for accounting noise in both the measurements *and* simulations in a principled manner using a so-called Bayesian errors-in-variables model (Gull, 2013; Minka, 1999). The noise related to the simulations can be due to segmentation errors and uncertainties in the tissue conductivity estimates. In general, we will refer to both errors and uncertainty as “noise” in the rest of the article. The Bayesian analysis was motivated as we found that the standard slope estimates were correlated with the measured field strengths, indicating that subjects where the signal is low had systematically worse regression fits (see Results section for details). In short, the regression model now becomes:

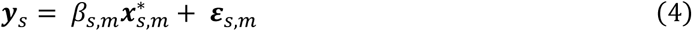

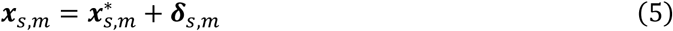

where 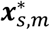 is a vector of the unobserved noiseless simulated values, and ***x***_*s,m*_ is a vector of the observed simulated potential differences. Note that in this model, both the measurements and the simulations are assumed to have noise. We further assume that the slopes are generated as:

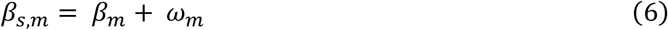

where *β*_*m*_is the unobserved group average of the slope for method *m*, and *ω*_*m*_is the noise, or rather the variation, of the slopes. All the noises and uncertainties are assumed to be Gaussian. We further need to define prior distributions on the noise parameters, the average slope *β*_*m*_, and the unobserved noiseless simulations 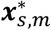 These, along with the full modeling details, can be found in the supplementary material. The Bayesian analysis was performed using Stan (Carpenter et al., 2017), specifically PyStan (https://pystan.readthedocs.io/en/latest/) for Python interfacing. The Stan code for running the analysis is provided in the supplementary material.

The benefit of adopting this type of Bayesian modeling is three-fold: first, as mentioned before, the noises in both the measurements and simulations can be estimated in a principled way. Second, we can study the posterior distributions of the slope for each subject and method to see which values are supported by the data given the model. Third, we can evaluate the group differences between the methods using the posterior predictive distribution of the slope for an unseen subject.

## RESULTS

### Experiment 1

Figure 1 shows the Dice scores comparing automated segmentations from the four pipelines to the manually corrected ones for the five main head tissue classes: WM, GM, CSF, skull and scalp. In general, the *SPM12*-based approaches, i.e., *headreco, headreco+CAT*, and *ROAST*, seem to perform on a somewhat similar level, whereas *mri2mesh* shows worse performance on all structures. The lower performance of *mri2mesh* is explained by two factors: first, as shown in Figure 2, the pipeline does not segment out the subcortical GM and segments the whole cerebellum as WM, resulting in lower Dice scores for WM and GM. Second, the extra-cerebral segmentations rely on a fairly simple method, which has been shown to be outperformed by the *SPM12*-based approaches (Nielsen et al., 2018). The largest differences between the two *headreco* pipelines and *ROAST* occur for WM and CSF. Here the reasons for the differences are more subtle, and likely explained by differences in the post-processing of the segmentations. The post-processing steps in the *headreco* pipelines seem to be somewhat more robust resulting in smaller spreads of the Dice scores.

**Figure 1:**
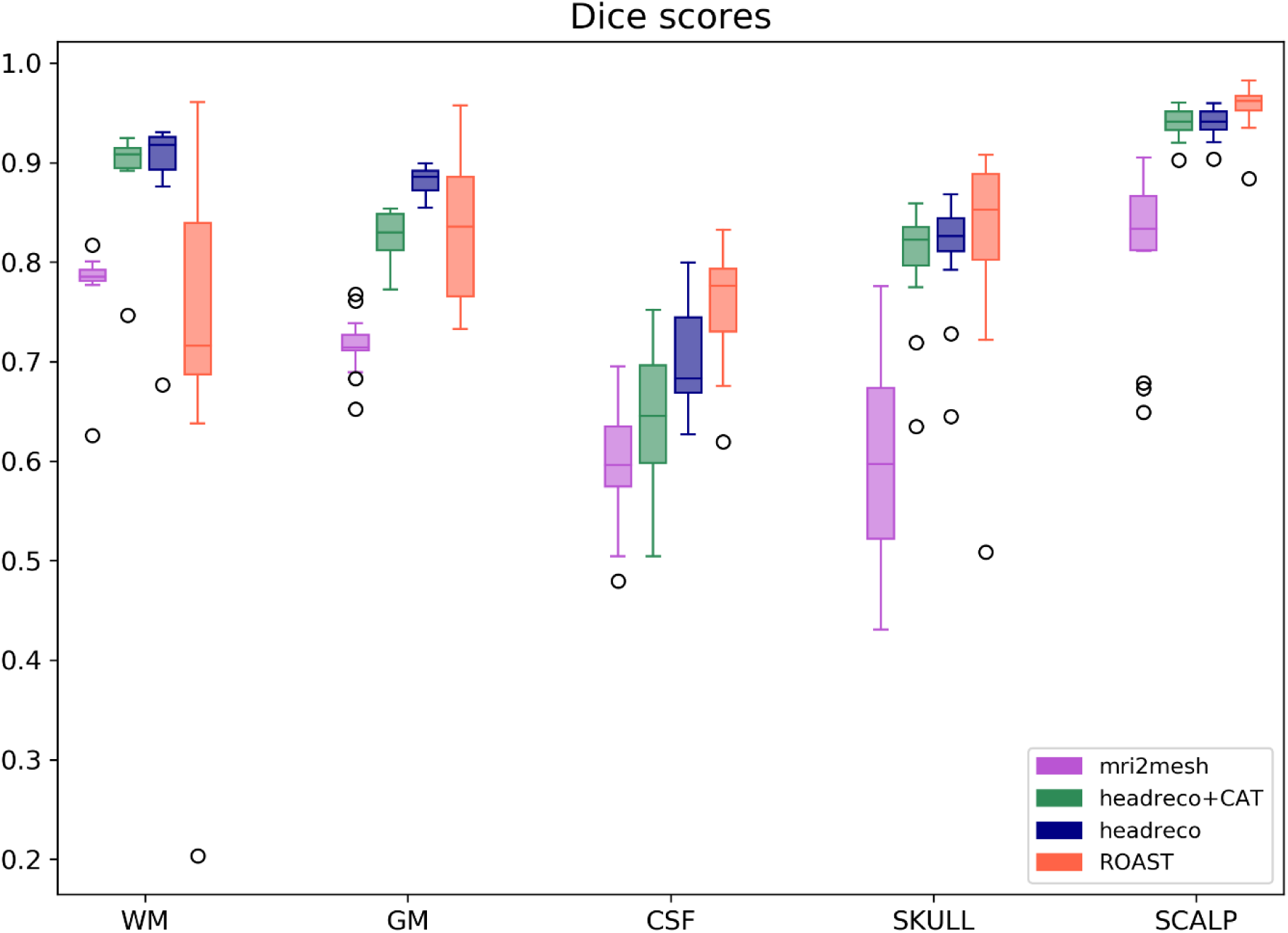
Dice scores computed for each of the pipelines in WM, GM, CSF, skull and scalp. On each box, the line marks the median, the box extends to lower and upper quartiles, whiskers extend up to 1.5 times the interquartile range, and data points beyond that are marked as outliers. The higher the score, more similar the automated segmentations are to the manually corrected ones.

Figure 2 shows two example segmentations from the data set. The first subject (P03) is a representative example of the quality of manually corrected and automated segmentations, whereas the second subject (P013) is a worst-case example of the manually corrected segmentations and corresponds to the WM outlier in the *ROAST* Dice scores. Note that the manually corrected segmentations suffer from general inaccuracies, such as the tendency to undersegment the sulcal CSF, and also include post-surgery corrections accounting for the craniotomy, intracranial electrodes and surgical drains. Thus, the Dice scores should not be interpreted as a direct measure of the segmentation accuracy, but rather highlighting general differences in the segmentations. For example, *headreco+CAT* seems to capture the GM sulci quite well but obtains a lower Dice score for both GM and CSF, which is due to the bias in the manually corrected segmentations. Example segmentations of all the subjects are provided in the supplementary material.

**Figure 2:**
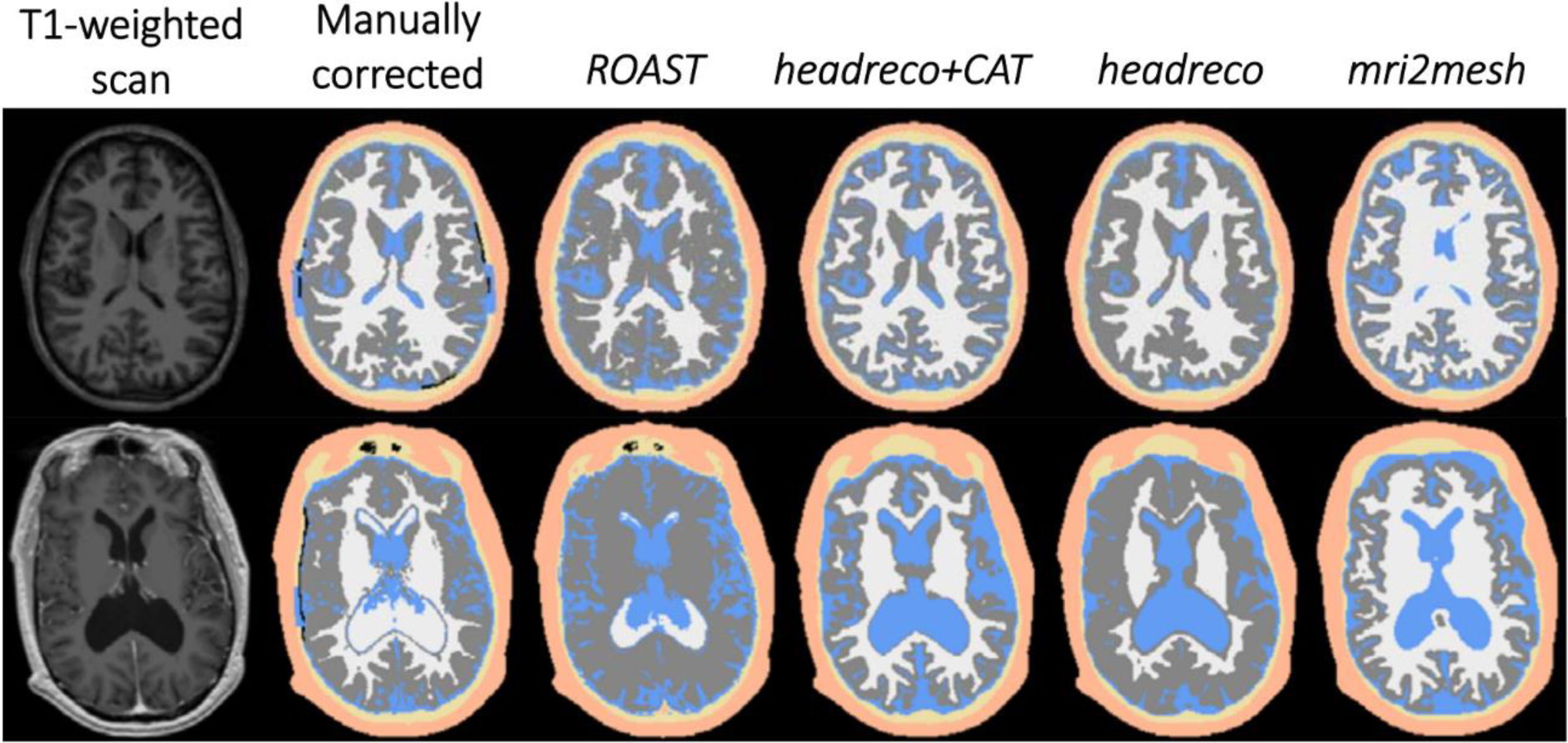
Example segmentations on subjects P03 (first row) and P013 (second row). From left to right: the input T1-weighted MRI scan, manually corrected segmentation, *ROAST, headreco+CAT, headreco* and *mri2mesh*. Note that *mri2mesh* does not segment the subcortical gray matter. The black lines close to the cortex in the manually corrected segmentations correspond to the intra-cranial electrode strips.

Figure 3 shows the mean of the relative difference between the electric fields (*eDiff, Eq. 2*) simulated using the electrode montages in the original study (Huang et al., 2017) for each pair of methods. The results from *headreco* and *headreco+CAT* are most similar to each other, which is expected as the only difference between the two is in the GM surface reconstruction. *mri2mesh* is the most different from all other methods, likely due to the different segmentation approach. *ROAST* presents moderate differences from the *headreco* methods. Both share large parts of the segmentation algorithm, as they are based on *SPM12*. However, post-processing of the segmentations and the electric fields differs, and is likely to cause most of the observed differences in the electric fields.

**Figure 3:**
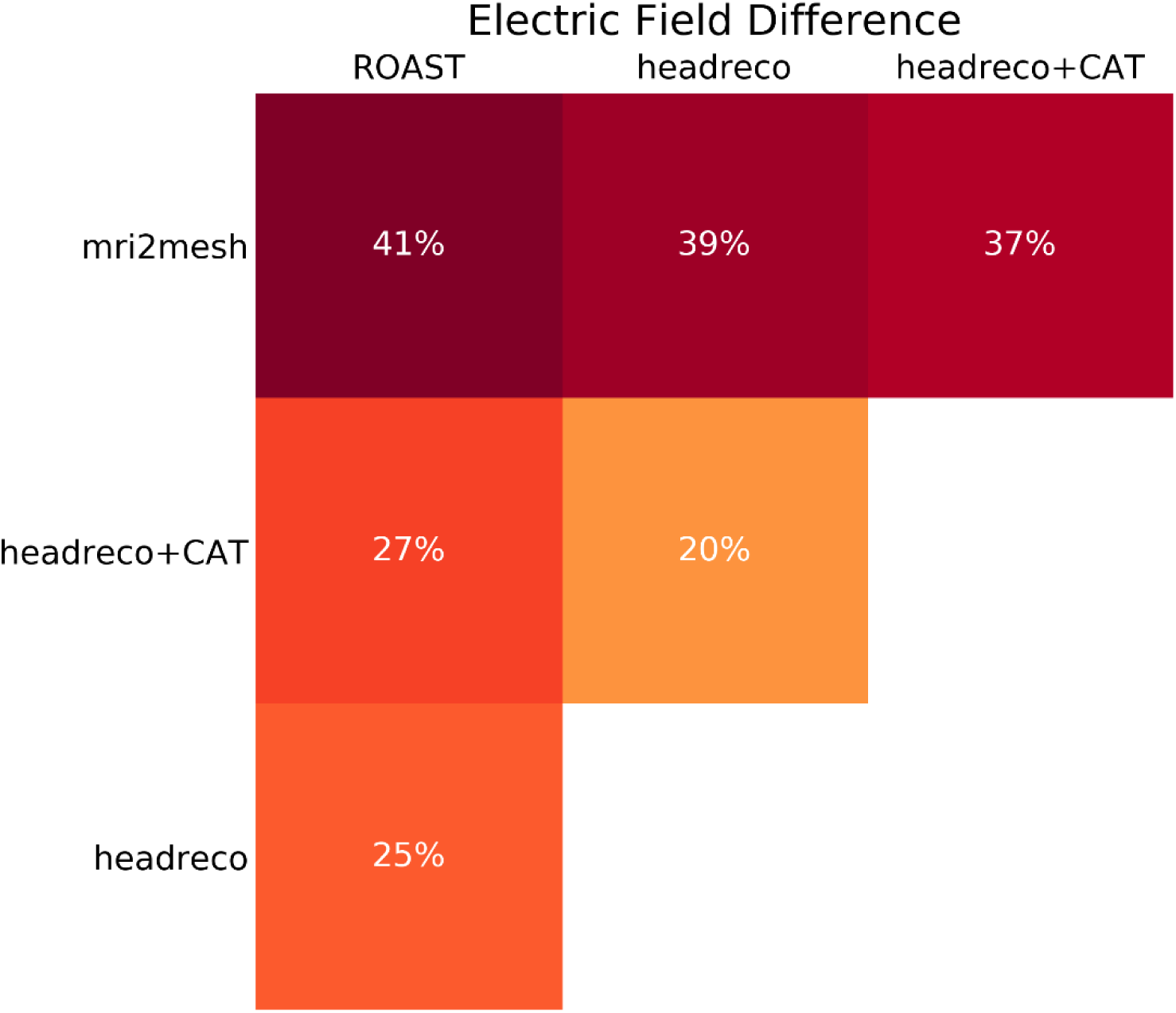
Mean electric field differences (*eDiff*) between the methods measured in overlapping gray matter.

In Figure 4, we show qualitative differences in the norm of the electric field stemming from differences in the anatomical segmentations. In general, we see that the amount of CSF has a large effect on the simulated electric fields likely due to shunting effects. The first row in Figure 4 shows that if the amount of CSF is less, the simulated fields in the cortex can be much higher as the current does not redistribute through the highly conducting CSF. Thus, accurate segmentation of the GM sulci also becomes important for locally accurate field modeling. The second row shows a similar effect, where the skull is mislabeled either as CSF (left) or scalp (right). Depending on where the skull border is placed, and the resulting thickness of the CSF layer, the estimated field strengths can be different. The final row in Figure 4, shows spurious islands of GM voxels in the *ROAST* segmentation, which can lead to extremely high field estimates in GM as these voxels are close to skull and surrounded by CSF. We note that segmenting out the CSF on this data set is challenging as no T2-weighted scan is provided. As the skull-CSF border is highly visible in T2 scans, they typically contribute to an accurate placement of the skull-CSF border (Nielsen et al., 2018). Also due to the lack of T2 data and the unreliability of the manual segmentations, it is hard to estimate which method has the most accurate reconstruction of the CSF.

**Figure 4:**
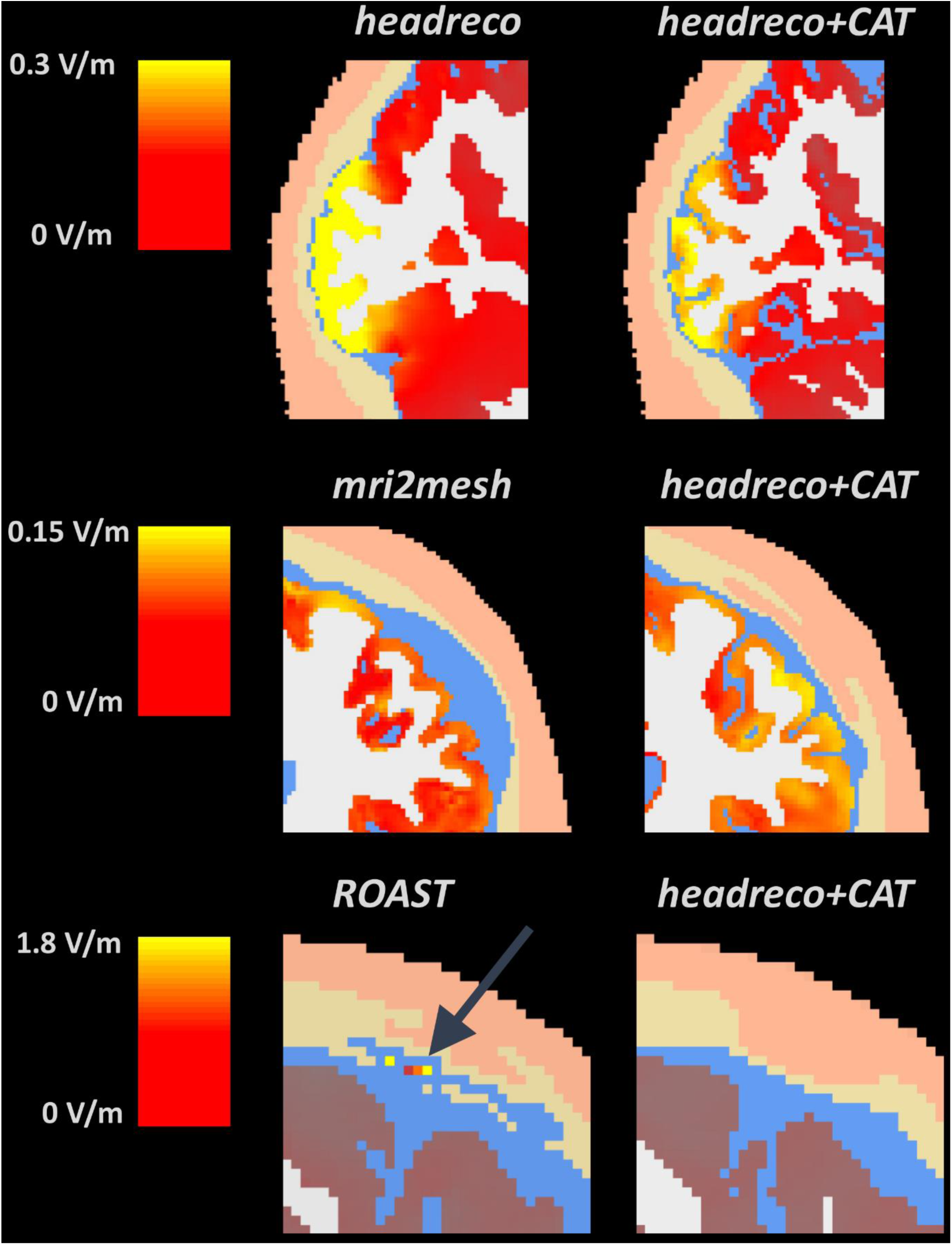
Examples of segmentation and electric field norm differences. First row: the amount of estimated CSF and differentiation of GM sulci result in different fields. Second row: erroneous segmentation of skull as CSF results in lower fields as the thicker CSF layer allows for more shunting. Third row: spurious islands of GM voxels in ROAST can have very large field estimates. Note that the electric field scale changes between the figure rows.

In Figure 5, we show the norm of the electric field in WM, GM and CSF for both *ROAST* and *headreco+CAT* in subject P03. The effect of the different electric field post-processings between *SimNIBS* and *ROAST* is quite striking: the interpolated field in *ROAST* is blurred, making the WM-GM border invisible and causing the large electric field estimates in the skull to bleed into CSF and to a lesser extent into GM. This effect makes the electric field estimates for CSF in *ROAST* clearly overestimated, as the fields in CSF are much lower due to its good electric conductivity. From Figure 5 it is also clear that the electric field differences reported in Figure 3 are very conservative as they were calculated only in the overlapping part of GM between the methods, discarding possibly large non-overlapping parts. However, we have opted for this approach due to the discontinuities of the electric field.

**Figure 5:**
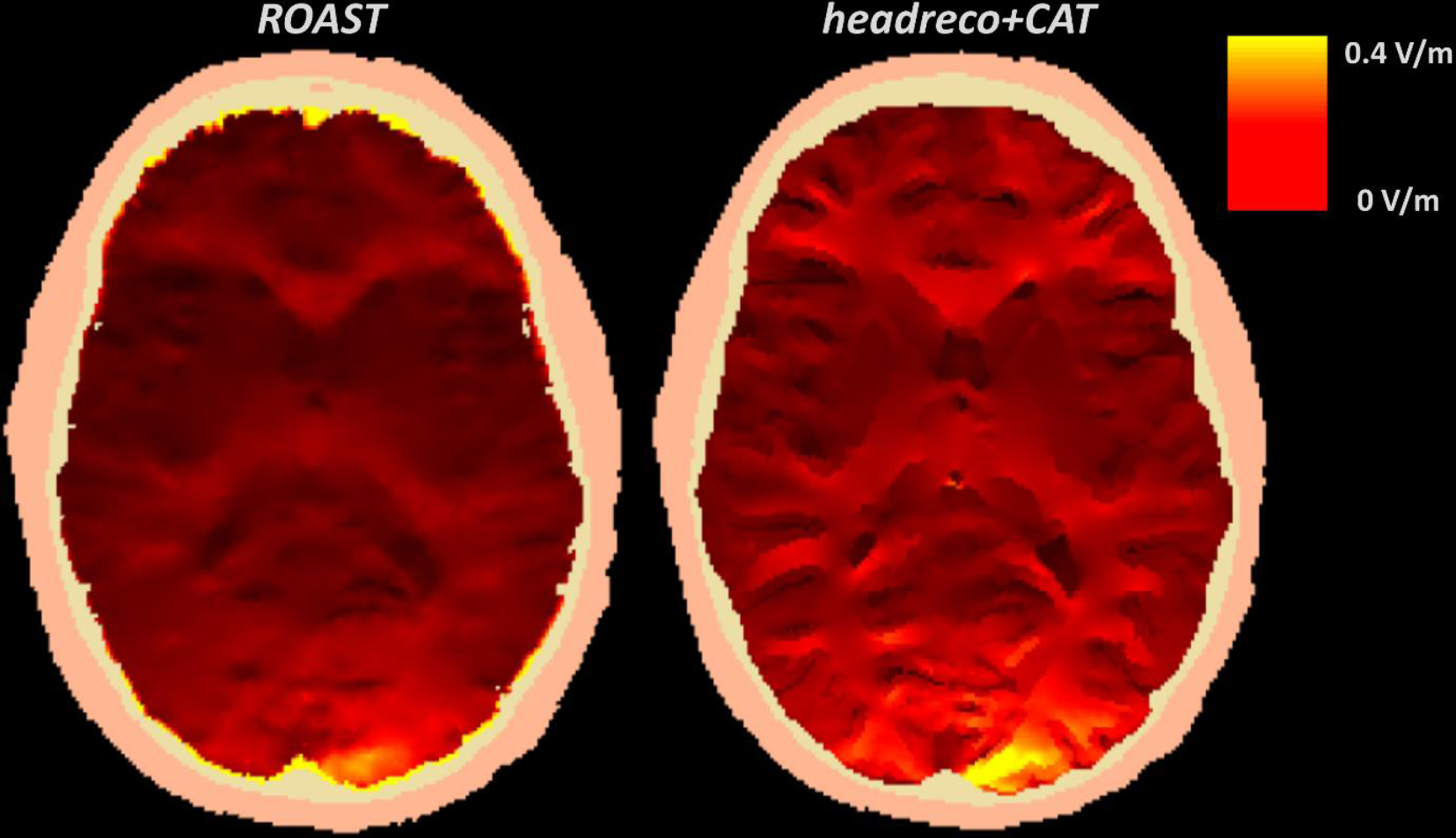
The distribution of the norm of the electric field over the full brain volume for example subject P03 from *ROAST* and *headreco+CAT*. Note the differences in smoothness of the simulated fields.

### Experiment 2

Figure 6 shows the coefficient of determination (r^2^) and the slope (*β*_*s,m*_) of the standard linear regression (Eq. 3) for each subject and method, and Table 1 shows the mean and standard deviation of both quantities, along with the correlation (*ρ*), across all subjects. Assessing Figure 6 qualitatively, it seems that all methods perform approximately equal in predicting the recordings in terms of both the r^2^and the slope. Testing for differences using a one-way repeated measures ANOVA revealed no statistically significant differences in the r^2^values between the methods (p=0.065) but did so for the slope values (p=0.021). However, pairwise post hoc comparisons between the slope estimates of methods did not reveal any differences using Tukey’s Honest Significant Difference test. When not considering multiple comparisons in the post hoc tests, we find trends of differences between the slope estimates of *headreco* and *headreco+CAT* (p=0.009), *headreco* and *ROAST* (p=0.017), and *headreco+CAT* and *mri2mesh* (p=0.027). The range of explained variance (r^2^) seems to be large over the subjects, where in some subjects (P04 and P07) the modeled fields explain the measurements well, while in others (P06 and P09) the prediction is poor. These results are in line with the ones reported in (Huang et al., 2018, 2017; Opitz et al., 2018). In addition, we also observe that all methods tend to overestimate the measured potential differences as reported by (Huang et al., 2018), but that the correlations are quite high for all methods and close to the ones reported for the head models generated from the manually corrected segmentations (Huang et al., 2017). However, in stark contrast to the comparisons in (Huang et al., 2018), we find no statistically significant differences in the accuracy of the field predictions between the methods.

**Table 1:** Mean *±* Standard deviation of the coefficient of determination (r^2^), slope (*β*) and correlation (*ρ*) for each head modeling pipeline, across all subjects.

|  | *mri2mesh* | *headreco+CAT* | *headreco* | *ROAST* |
| --- | --- | --- | --- | --- |
| Coefficient of Determination ($r^2$) | $0.506 \pm 0.163$ | $0.580 \pm 0.150$ | $0.567 \pm 0.148$ | $0.554 \pm 0.175$ |
| Slope ($\beta$) | $0.526 \pm 0.284$ | $0.632 \pm 0.313$ | $0.556 \pm 0.247$ | $0.644 \pm 0.353$ |
| Correlation ($\rho$) | $0.700 \pm 0.127$ | $0.755 \pm 0.106$ | $0.746 \pm 0.103$ | $0.734 \pm 0.128$ |

**Figure 6:**
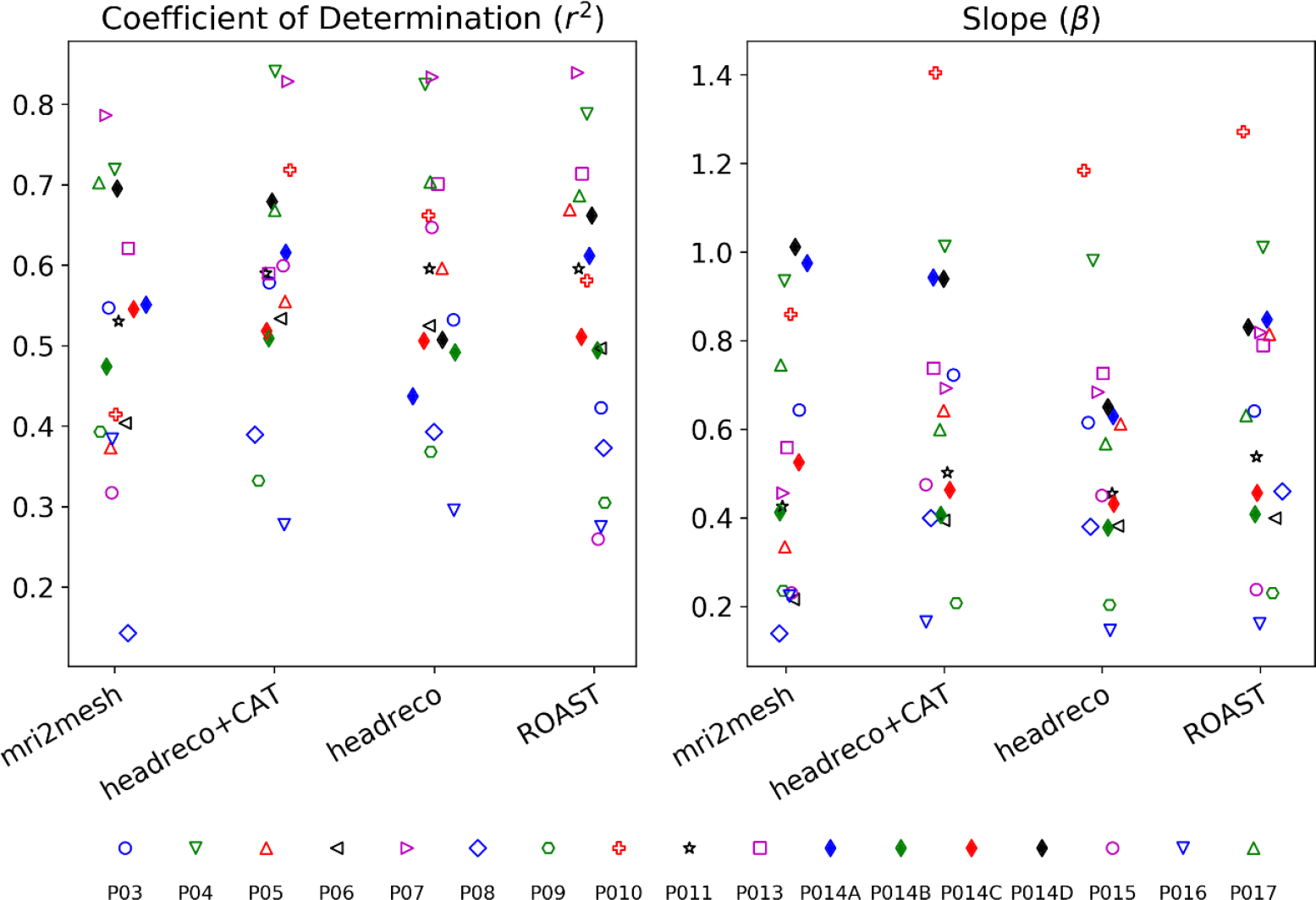
Coefficient of determination (r^2^) and slope (*β*_*s,m*_) of the linear fit, for all subjects and methods.

Figure 7 shows the correlation between the strength of the recorded potential differences and the slope estimates along with p-values for the correlation. We find a statistically significant correlation for all methods except *headreco*, which implies that the slope estimates are likely biased due to noise present in both the measurements and the simulations. In fact, it is well-known that if noise in the so-called independent, or predictor, variables is unaccounted for, the regression coefficient will be underestimated (Frost and Thompson, 2000; Fuller, 1987). The problem persists even if the predicted and independent variables are exchanged as then the noise in the measurements is ignored, or if an intercept term is added. This prompted us to conduct a full Bayesian regression analysis (Eq. 4-5), as outlined in the Experiments and detailed in the supplementary material.

**Figure 7:**
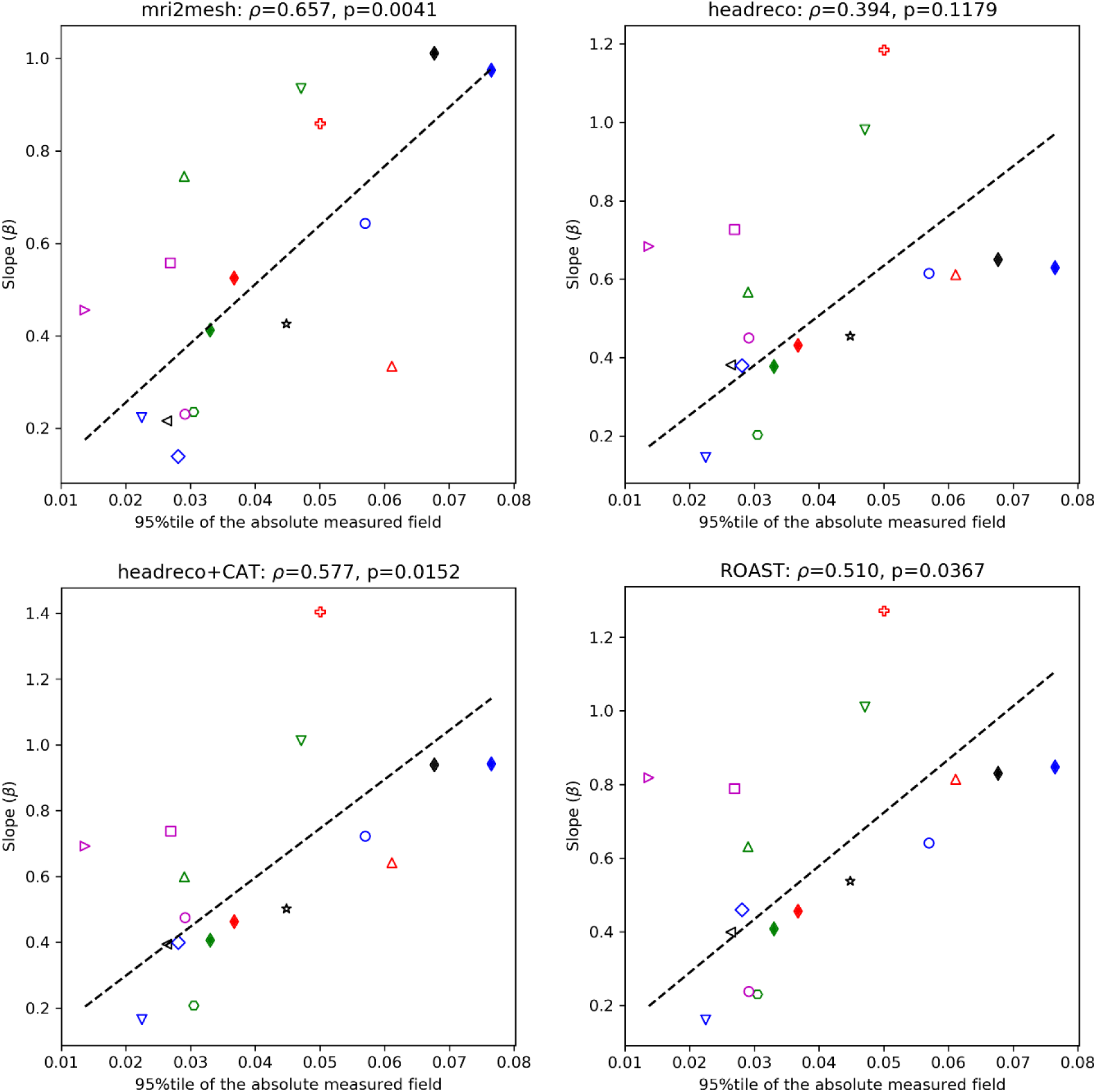
Correlation between the 95^th^ percentile of the absolute measured field and the slope estimates. Linear regression line without intercept shown in black. The correlation is significant for all methods except *headreco*.

To demonstrate how the regression estimates are linked to the range of the measured and simulated fields and therefore, indirectly, to the choice of electrode positioning, we look at subject P014. This subject is interesting as the locations of the stimulation electrodes were varied during the experiment. Results in Figure 6 show that configurations P014A and P014D seem to be better fitted than P014B and P014C for all methods apart from *headreco* where the difference is not as clear. Figure 8 shows the results of the Bayesian analysis for configurations P014C and P014D. The measured potential differences in the former are closer to zero and look relatively noisier than in the latter. This results in a lower slope estimate and a wider 95% compatibility interval for P014C compared to P014D. The results suggest that some of the poor fits observed in Figure 6 are due to the signal being drowned by noise, in measurements and simulations, which would also explain the correlation of the signal strength with the slope shown in Figure 7. Detailed analysis, similar to Figure 8, on all subjects can be found in the supplementary material.

**Figure 8:**
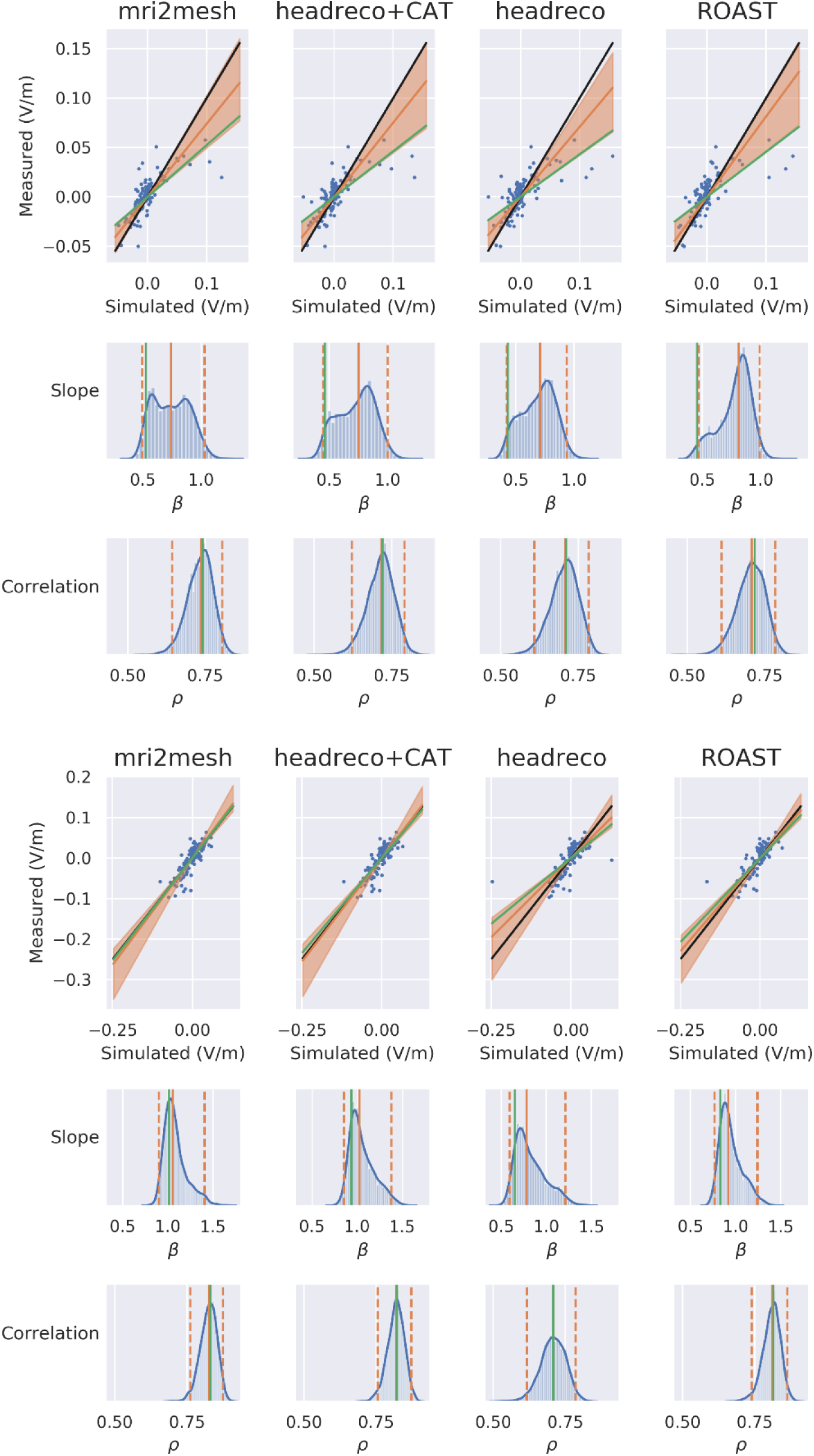
Bayesian regression analysis for subject P014. The first panel shows results for P014C, and the second panel for P014D. In each scatter plot the black line denotes a slope of one, the orange line is the median of the posterior distribution for the slope with the shading denoting 95% compatibility interval, and the green line is the standard slope fit (as in Figure 6). The histograms show the posterior distributions of the slope and correlation with the median denoted as an orange solid line, the 95% interval as a dashed line, and the green solid line denoting the standard fit (as in Figure 6). Similar plots for all subjects are included in the supplementary material.

As seen in Figure 8 the slopes from the standard analysis seem to be underestimated compared to the median of the posterior distribution obtained from the Bayesian analysis. This is in general true for most subjects, as shown in the supplementary material. To quantify the underestimation, we sampled 4000 slope estimates from the normal distribution governing the slope of the standard regression in each subject, compared those in a pairwise manner to the slope samples from the posterior distribution in each subject, and computed the probability that the standard regression slope is smaller than the corresponding Bayesian one. The probabilities computed this way are: 0.867 for *mri2mesh*, 0.866 for *headreco+CAT*, 0.869 for *headreco*, and 0.861 for *ROAST*. The distribution of these pooled differences for each method over all subjects is shown in Figure 9. All of the distributions have medians larger than zero and the area to the right from zero corresponds to the probabilities stated above. To link this to a more classical statistical analysis, we performed paired Wilcoxon signed-rank tests between the standard slope estimates (Figure 6) and the posterior means, which resulted in the p-values < 0.001 for all segmentation methods. Thus both the Bayesian analysis and a standard pairwise test between the slope estimates indicate that the regression results in Figure 6 and Table 1, along with the results in (Huang et al., 2018) and (Huang et al., 2017) are underestimating the true slopes as they do not account for the noise in the simulation results. The analysis, results, and conclusions presented in (Huang et al., 2018) and (Huang et al., 2017) would benefit from being revisited with this underestimation in mind. In Figure 10 we show the pooled difference plots between correlation samples from the standard analysis (Figure 6, left and Table 1, second row), which were first Fisher transformed and then assumed normally distributed (Fisher, 1921, 1915), and the corresponding Bayesian alternatives. All of the distributions are centered on zero meaning that in general the difference between the standard and Bayesian correlation samples is small, although the 95% compatibility interval ranges from approximately −0.2 to 0.2. These results seem to be in line with those presented in Figure 6 for r^2^, indicating that the correlation estimates from the standard and Bayesian analysis agree to a large extent on this data set.

**Figure 9:**
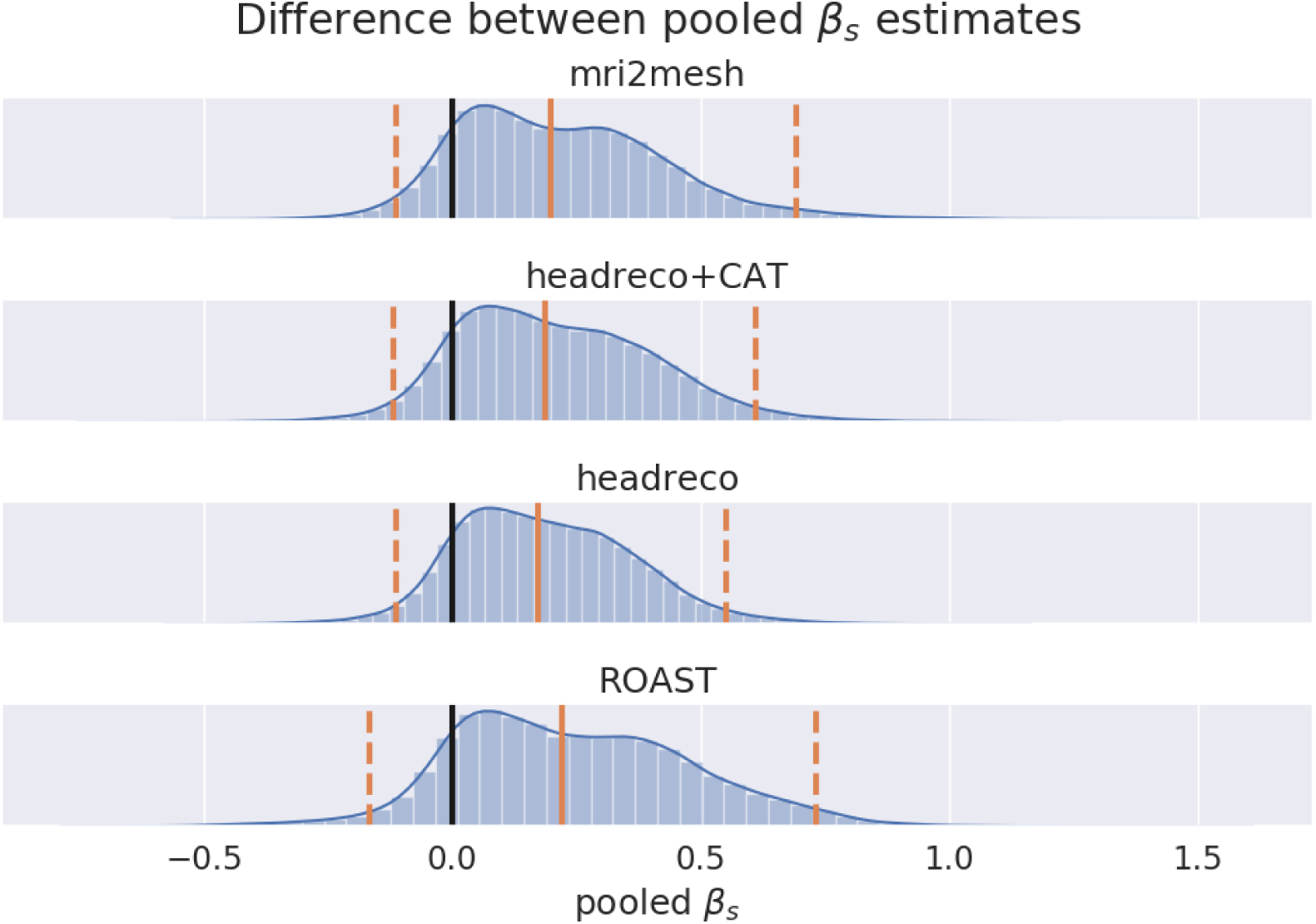
Distribution of the differences between the slope samples from the standard and Bayesian analysis pooled over all subjects. Solid orange line denotes the median, dashed lines the 95% compatibility interval and the black line at zero indicates no difference. The distributions are shifted towards the right indicating a difference between the standard and Bayesian slope estimates.

**Figure 10:**
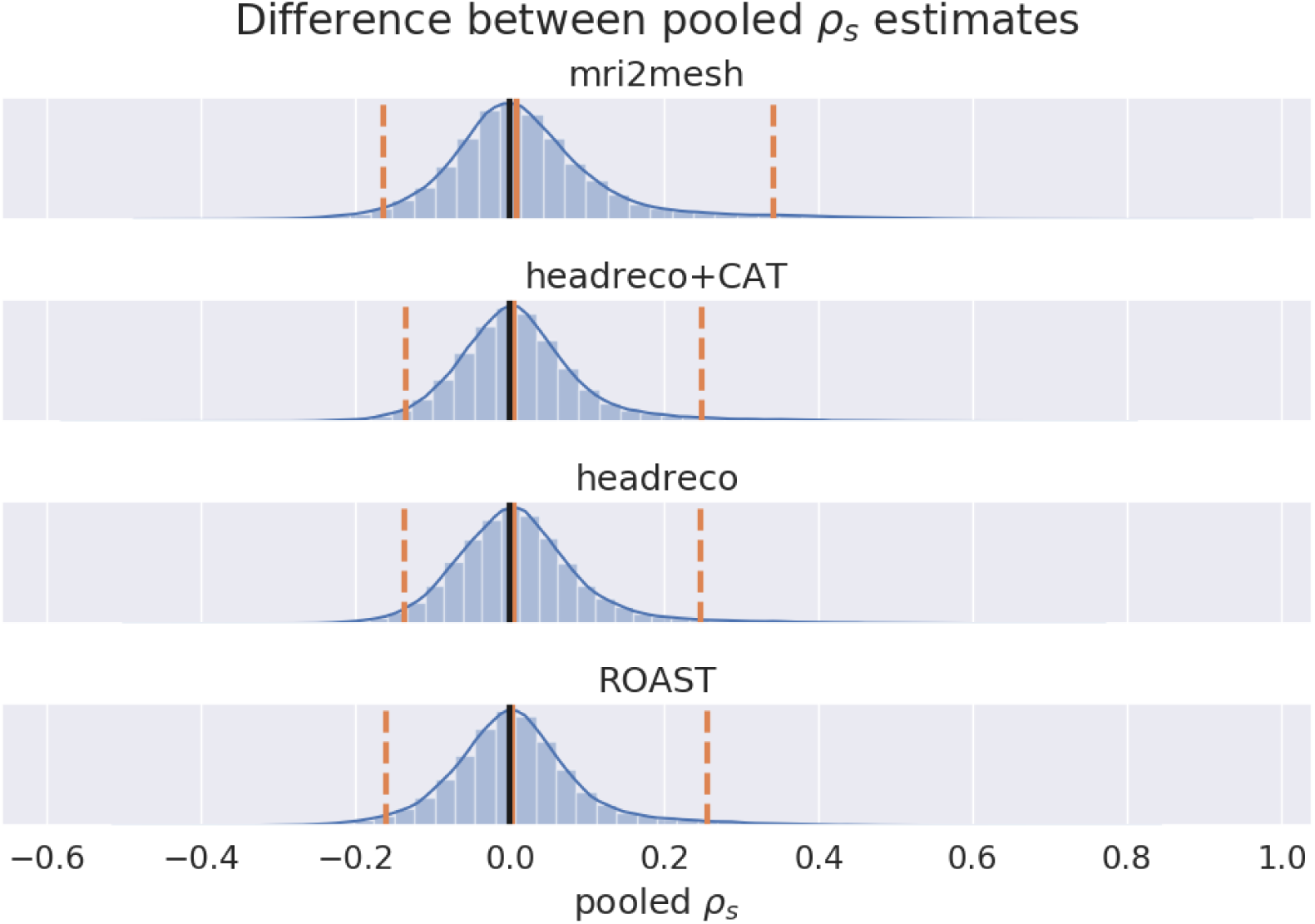
Distribution of the differences between the regression samples from the standard regression and the samples from the posterior distribution pooled over all subjects. Solid orange line denotes the median, dashed lines the 95% compatibility interval and the black line at zero indicates no difference. The distributions are centered on zero indicating that, in general, the differences between the standard and Bayesian correlation estimates are small.

To compare differences between the methods, we pooled the differences between the slope samples from the posterior distributions for each pair of methods over all subjects. The distribution is shown in Figure 11, and the individual slope posteriors for all subjects and methods are plotted in the supplementary material. In general we find that the differences between methods are small as the peaks of the distributions are close to zero, although more extreme differences are also supported by the model given the data. The pooled difference distributions for the correlations between the methods are shown in Figure 12. The distributions are all centered on zero indicating small differences in the correlation estimates between methods but are heavy tailed also supporting larger differences. Finally, Figure 13 shows the posterior predictive distributions of the slope 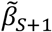 for an unseen subject given the data, overlaid with the median and 95% confidence interval of the slopes estimated using the standard regression analysis (Figure 6). The posterior predictive distribution tells us, which slope values we should expect, given the data we have seen, if a new subject were to be measured. The posterior predictive distributions of all methods cover a large range of possible slope values as a result of the high variability in the recorded data and simulations. Table 2 lists summary statistics of the predictive distributions for each method, along with the summary statistics of the correlation distributions pooled over subjects. As mentioned above, the correlation estimates from the standard analysis match the ones obtained from the Bayesian analysis, while the slopes from the standard analysis are systematically lower, but both analyses reveal no large differences between the methods on this data set.

**Table 2:** Posterior median, 97.5^th^ percentile (+) and 2.5^th^ percentile (-) of the slope 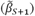 and correlation 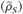 for each head modeling pipeline, across all subjects. The values correspond to the orange lines in Figures 10 and 11.

|  | *mri2mesh* | *headreco+CAT* | *headreco* | *ROAST* |
| --- | --- | --- | --- | --- |
| Slope ($\tilde{\beta}_{S+1}$) | $0.749 \pm_{0.202}^{1.276}$ | $0.841 \pm_{0.196}^{1.438}$ | $0.741 \pm_{0.254}^{1.257}$ | $0.862 \pm_{0.402}^{1.282}$ |
| Correlation ($\tilde{\rho}_S$) | $0.704 \pm_{0.341}^{0.891}$ | $0.759 \pm_{0.434}^{0.926}$ | $0.740 \pm_{0.448}^{0.923}$ | $0.738 \pm_{0.365}^{0.920}$ |

**Figure 11:**
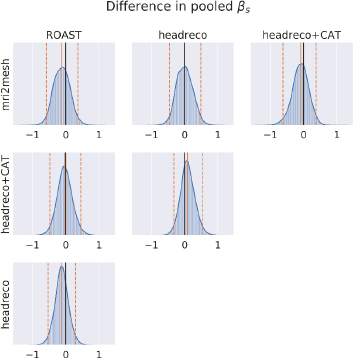
Distributions of the pairwise differences (row method – column method) between the samples from the posterior distribution of the slope pooled over subjects. Solid orange line denotes the median, dashed lines the 95% compatibility interval, and the solid black line at zero indicates no difference.

**Figure 12:**
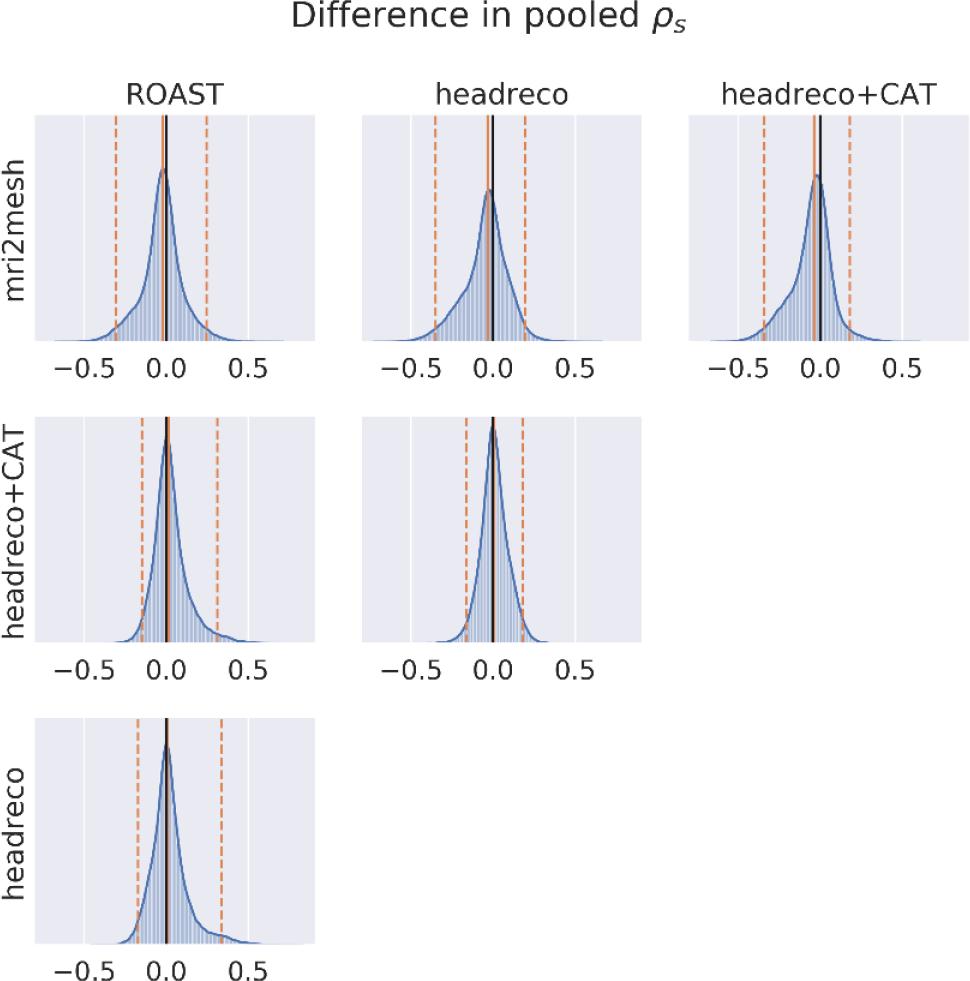
Distributions of the pairwise differences (row method – column method) between the samples from the posterior distribution of the correlation pooled over subjects. Solid orange line denotes the median, dashed lines the 95% compatibility interval, and the solid black line at zero indicates no difference.

**Figure 13:**
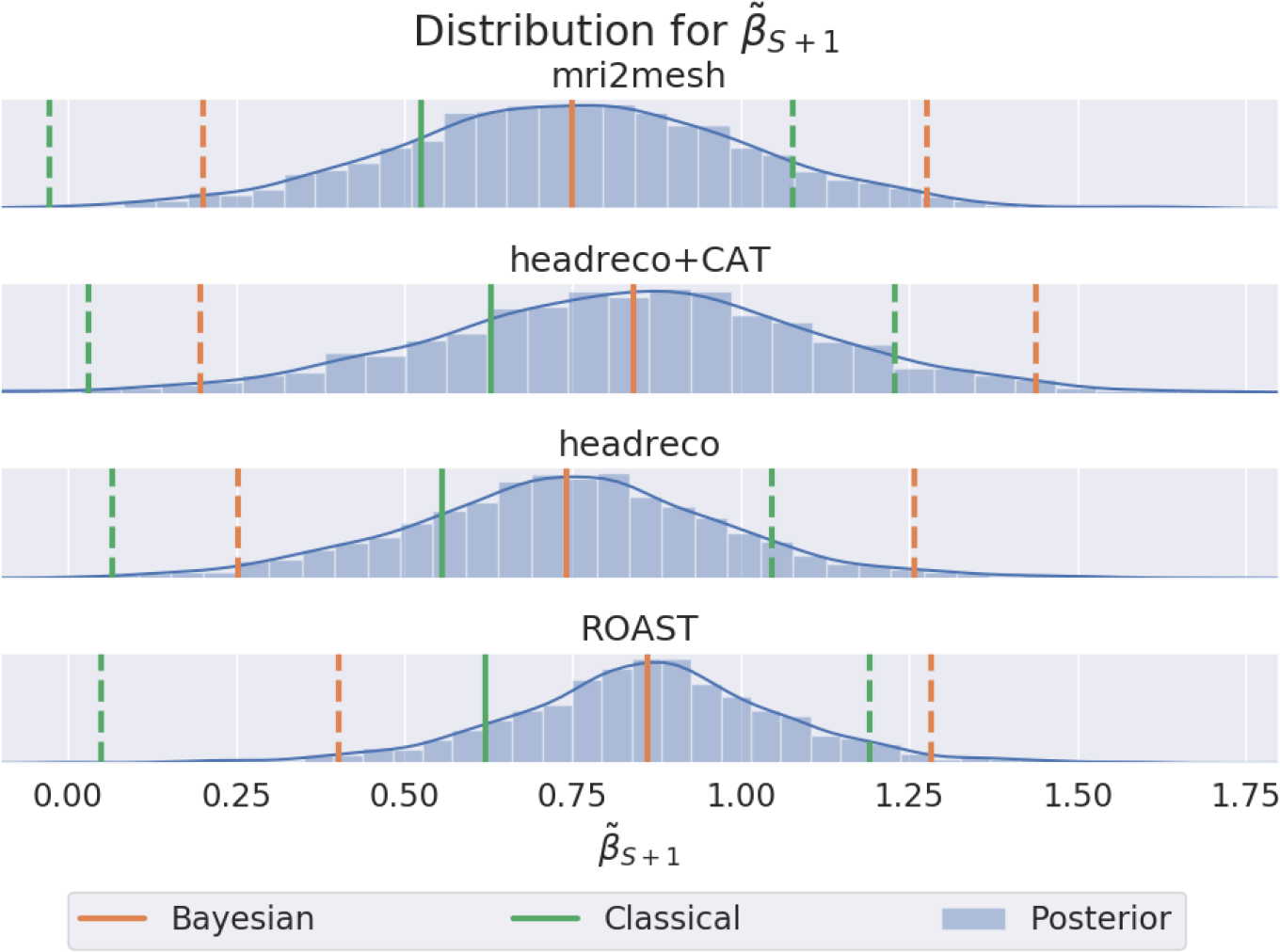
Posterior predictive distribution of the slope if a new subject were to be measured. Solid orange line denotes the median and dashed lines the 95% compatibility interval. The corresponding green lines denote the estimates for the median and 95% confidence interval for the standard analysis as in Figure 6.

## DISCUSSION

In the current work, we used an openly available dataset (Huang et al., 2017) with intracranial electric potential recordings, MR scans and manually corrected segmentations to compare four segmentation methods coming from two packages for TES electric field simulations: SimNIBS 2.1 (Saturnino et al., 2018) and ROAST 2.7 (Huang et al., 2018). We started by comparing the segmentations, where we found large differences between the methods. However, we also found the manually corrected segmentations in the dataset to be not accurate in all subjects. Next, we compared electric fields in gray matter calculated using the different software on the different subjects. We found there to be large differences (up to 41%) between the segmentation methods and packages. These seem to come mostly from differences in the MR segmentation and post-processing methods during head meshing, though differences in the post-processing of the calculated electric potentials also contribute. Afterwards, we compared the recorded and simulated potential differences at the electrodes using a standard linear regression analysis and a Bayesian errors-in-variables model. There, we observed no large differences between the segmentation methods in both types of analyses. Additionally, we find that the standard linear regression, by not considering possible errors in the simulations, underestimates the slope of the linear relationship between recordings and simulations.

The results highlight two important points: first, the noise in both the electric field simulations and measurements needs to be considered in the analysis in order to obtain reliable regression estimates, and second, even though the electric field simulations differ between the methods this difference does not show up in the correlation with the recordings. In intra-cranial electrode measurements, the locations of the recording electrodes are decided solely based on clinical considerations, and thus they might not be in a region experiencing the highest electric field strengths for the given stimulation protocol. As exemplified by subject P014 in Figure 8, when the recorded signal is low, noise in the measurements and simulations leads to regression estimates with a high variance as the data does not have enough information to pin down the parameter values. This observation suggests the need for a more strategic approach to head model validation, opting for tailored stimulation protocols in each subject to produce enough signal so that conclusive validation is possible. This is crucial as the intracranial electrodes can measure potential differences only in the plane defined by the electrode strip, meaning that changes in the electric field perpendicular to the measurement plane are not picked up (Huang et al., 2017). Measurements of the electric field in all three directions would likely distinguish differences between field simulations better but is difficult in practice with intra-cranial electrodes. Complementing intra-cranial recordings with new, non-invasive techniques, such as MRCDI, for volume measurements of the electric fields seems thus important for future validations of individualized field modeling.

Likely due to restrictions imposed by studying a clinical population, only T1-weighted MR scans were included in the dataset and the quality of the images is variable. The latter is to an extent expected, as clinical scans typically include more artifacts related to movement. The scans in this data set seem to have a fairly high resolution, although the specific scan parameters and original resolution are not reported in (Huang et al., 2017). The lower parts of the heads are missing, rendering the anatomical accuracy of the corresponding parts of the head volume conductor models uncertain. In some subjects, we also observed reduced WM-GM contrast (see Figure 2, second row) and resolutions lower than 1mm^3^. The quality of the MR scans will certainly influence the anatomical accuracy of the generated head models, increasing noise in the resulting simulations and making comparisons between methods harder. In particular, automatic methods so far achieve robust segmentations of the skull compartment only when the T1-weighted image is complemented by additional images with good contrast between CSF and skull, such as T2-weighted scans (Nielsen et al., 2018). The manually corrected segmentations also suffer from some inaccuracies, which makes it difficult to establish a baseline between simulations and measurements, and to link anatomical accuracy to simulation results. That being said, this open-source data set is at the moment still unique and we acknowledge its importance for validating volume conductor models of the head. We would like to encourage its further curation to ameliorate some of the mentioned limitations.

This data set has been analyzed in a previous study (Huang et al., 2018) to compare the electric field simulations from the same toolboxes used in this article. The authors find that *headreco* and *headreco+CAT* have significantly lower correlations with the measurements and linear slopes than *ROAST*. The difference seems to be driven by outliers where in some subjects the electric potentials are zero for *headreco* and *headreco+CAT* (Huang et al., 2018). In this article, when repeating the analysis (Figure 6 & Table 1) we found no differences between the methods and observe no outliers for any of the methods. We encourage users of simulation software to always check the segmentation and simulations results for obvious errors. Following this procedure, we changed some of the default values for specific subjects, as detailed in the Experiments section, so that sensible simulations were obtained for all methods.

In prior articles based on this dataset (Huang et al., 2018, 2017), the comparisons between simulations and recordings rely on a standard linear regression analysis where the simulations are assumed be noiseless. Thus, conclusions that the simulations overestimate the electric fields in the brain, which are based on the slope of the regression results, are not necessarily warranted and are hard to put into perspective as only point estimates are provided. This ties together with the optimization of tissue conductivities in (Huang et al., 2017), which relies on minimizing the squared error between the simulations and recording, with both being assumed noiseless. It seems likely that this procedure provides biased conductivity estimates. Given that the validation of electric field simulations is an important issue to tackle in developing TES into a clinically relevant tool, our results suggest the need for a careful interpretation of the existing data based on suited analyses methods. We also hope that the issues raised here give helpful guidance for planning future validation attempts.

## ACKNOWLEDGEMENTS

We would like to thank Hartwig R. Siebner for helpful comments on the manuscript. This study was supported by the Lundbeck foundation (grant no. R244-2017-196), the Novonordisk foundation (grant no. NNF14OC0011413).

This project has received funding from the European Union’s Horizon 2020 research and innovation programme under grant agreement No. 731827 (STIPED).

