## Supplementary 1 for "Comparing and Validating Automated Tools for Individualized Electric Field Simulations in the Human Head"

April 17, 2019

### S.1 Bayesian errors-in-variables regression

The Bayesian errors-in-variables linear regression model used in the article closely follows the treatment presented in [1]. We start by defining the likelihood model for the observed voltage recordings $\mathbf{y}_s = (y_{s,1}, \ldots, y_{s,N})^T$ and simulation results $\mathbf{x}_{s,m} = (x_{s,m,1}, \ldots, x_{s,m,N})^T$. Here, $s$ denotes subject, $m$ denotes one of the four different simulation methods, and $N$ is the number of measurements. The likelihood model is assumed to be a multivariate normal:

$$p\left(\mathbf{y}_s, \mathbf{x}_{s,m} | \boldsymbol{\mu}_{s,m,n}, \boldsymbol{\Sigma}_{s,m}\right) = \prod_{n=1}^{N} p\left((y_{s,n}, x_{s,m,n}) | \boldsymbol{\mu}_{s,m,n}, \boldsymbol{\Sigma}_{s,m}\right) \tag{S.1}$$

$$= \prod_{n=1}^{N} \mathcal{N}\left((y_{s,n}, x_{s,m,n}) | \boldsymbol{\mu}_{s,m,n}, \boldsymbol{\Sigma}_{s,m}\right), \tag{S.2}$$

where we assumed that each pair $n$ of recordings and simulations are drawn independently. The mean vector is defined as: $\boldsymbol{\mu}_{s,m,n} = (\beta_{s,m}, 1)x^*_{s,m,n}$ where $x^*_{s,m,n}$ denotes the unobserved noiseless simulation and $\beta_{s,m}$ is the slope term of the linear regression. The noise covariance matrix is defined as:

$$\mathbf{\Sigma}_{s,m} = \begin{bmatrix} \sigma^2_{y,s} & 0 \\ 0 & \sigma^2_{x,s,m} \end{bmatrix}, \tag{S.3}$$

where $\sigma^2_{y,s}$ and $\sigma^2_{x,s,m}$ are the noise variances of the recordings and simulations respectively. Note that the observed values are conditionally independent of each other given the unobserved noiseless simulation parameter $x^*_{s,m,n}$.

Next we define the distributions on the unobserved simulation parameter and the regression slope:

$$p(x^*_{s,m,n}|\sigma_{t,s,m}) = \mathcal{N}(x^*_{s,m,n}|0, \sigma^2_{t,s,m}) \tag{S.4}$$
$$p(\beta_{s,m}|\beta_m, \sigma_{\beta,m}) = \mathcal{N}(\beta_{s,m}|\beta_m, \sigma^2_{\beta,m}), \tag{S.5}$$

where $\sigma^2_{t,s,m}$ is the variance of the unobserved simulation parameters, $\beta_m$ is the group average slope for method $m$, and $\sigma^2_{\beta,m}$ the slope variation.

Finally, we define prior distributions on the means and the scales of the distributions:

$$p(\beta_m) = \mathcal{N}(\beta_m|0, 4), \tag{S.6}$$
$$p(\sigma_{\beta,m}) = \mathcal{C}^+(\sigma_{\beta,m}|0, 1), \tag{S.7}$$
$$p(\sigma_{t,s,m}) = \mathcal{C}^+(\sigma_{t,s,m}|0, 0.2), \tag{S.8}$$
$$p(\sigma_{x,s,m}) = \mathcal{C}^+(\sigma_{x,s,m}|0, 0.2), \tag{S.9}$$
$$p(\sigma_{y,s}) = \mathcal{C}^+(\sigma_{y,s}|0, 0.2) \tag{S.10}$$

where $\mathcal{C}^+(\cdot)$ is a half-Cauchy distribution. The prior distributions are shown in Figure S.1.

The Stan toolbox relies on sampling to do the inference in the joint distribution defined by the model above. To limit the number of parameters, we marginalize over the unobserved noiseless simulation parameter. Following [1] this becomes:

$$\begin{aligned} &p\left((y_{s,n}, x_{s,m,n})|\beta_{s,m}, \sigma_{y,s}, \sigma_{x,s,m}, \sigma_{t,s,m}\right) = \\ &\int_{x^*_{s,m,n}} p\left((y_{s,n}, x_{s,m,n}), x^*_{s,m,n}|\beta_{s,m}, \sigma_{y,s}, \sigma_{x,s,m}, \sigma_{t,s,m}\right) dx^*_{s,m,n}, \end{aligned} \tag{S.11}$$

where

$$\begin{aligned} &p\left((y_{s,n}, x_{s,m,n}), x^*_{s,m,n}|\beta_{s,m}, \sigma_{y,s}, \sigma_{x,s,m}, \sigma_{t,s,m}\right) = \\ &\quad p\left((y_{s,n}, x_{s,m,n})|\mu_{s,m,n}, \mathbf{\Sigma}_{s,m}\right) p(x^*_{s,m,n}|\sigma_{t,s,m}). \end{aligned} \tag{S.12}$$

The joint distribution of the observations and unobserved simulation variable is a product of two Gaussian distributions, so the resulting distribution is also Gaussian.

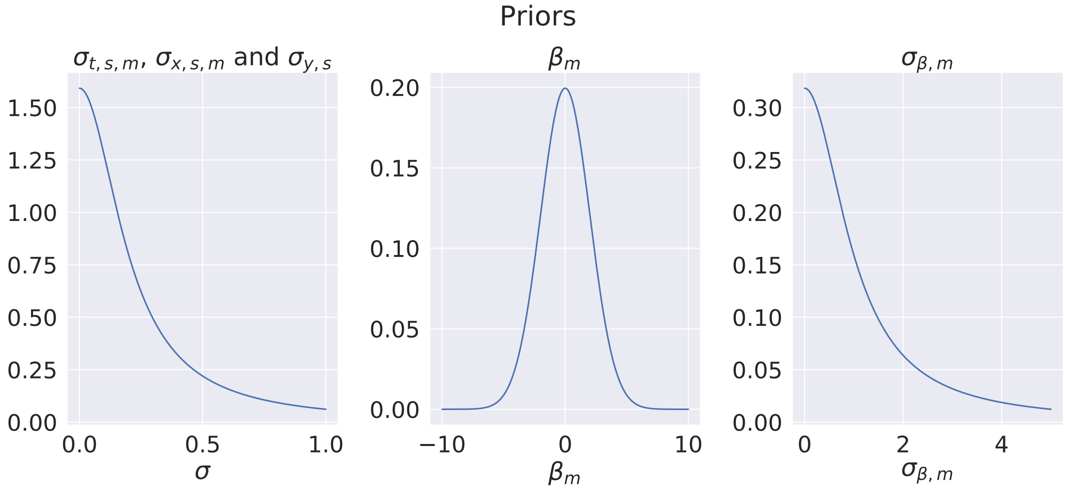


Figure S.1: Prior distributions

The integration yields [1]:

$$p\left((y_{s,n}, x_{s,m,n})|\beta_{s,m}, \sigma_{y,s}, \sigma_{x,s,m}, \sigma_{t,s,m}\right) = \mathcal{N}\left((y_{s,n}, x_{s,m,n})|\mathbf{0}, \sigma_{t,s,m}^2 \mathbf{\Sigma}_{\beta,s,m} + \mathbf{\Sigma}_{s,m}\right), \tag{S.13}$$

where

$$\mathbf{\Sigma}_{\beta,s,m} = \begin{bmatrix} \beta_{s,m}^2 & \beta_{s,m} \\ \beta_{s,m} & 1 \end{bmatrix}. \tag{S.14}$$

Finally the marginalized likelihood becomes:

$$p\left(\mathbf{y}_s, \mathbf{x}_{s,m}|\sigma_{t,s,m}, \mathbf{\Sigma}_{\beta,s,m}, \mathbf{\Sigma}_{s,m}\right) = \prod_{n=1}^{N} p\left((y_{s,n}, x_{s,m,n})|\sigma_{t,s,m}, \mathbf{\Sigma}_{\beta,s,m}, \mathbf{\Sigma}_{s,m}\right) \tag{S.15}$$

$$= \prod_{n=1}^{N} \mathcal{N}\left((y_{s,n}, x_{s,m,n})|\mathbf{0}, \sigma_{t,s,m}^2 \mathbf{\Sigma}_{\beta,s,m} + \mathbf{\Sigma}_{s,m}\right). \tag{S.16}$$

Note that the observation are now dependent through $\mathbf{\Sigma}_{\beta,s,m}$. The posterior distribution over the model parameters can now be written as:

$$p(\beta_{s,m}, \beta_m, \sigma_{\beta,m}, \sigma_{t,s,m}, \sigma_{x,s,m}, \sigma_{y,s}|\mathbf{y}_s, \mathbf{x}_{s,m}) \propto$$
$$p\left(\mathbf{y}_s, \mathbf{x}_{s,m}|\sigma_{t,s,m}, \mathbf{\Sigma}_{\beta,s,m}, \mathbf{\Sigma}_{s,m}\right) p(\beta_{s,m}|\beta_m, \sigma_{\beta,m}) p(\beta_m) p(\sigma_{\beta,m}) p(\sigma_{t,s,m}) p(\sigma_{x,s,m}) p(\sigma_{y,s}) \tag{S.17}$$

Listing 1: Stan code for sampling from the model

```
data {
    int N; // Total number of recordings in each method
    int S; // Number of subjects
    vector[N] x;
    vector[N] y;
    int n_dat[S + 1];
    // Priors for scale parameters (Cauchy)
    real loc_sigma_prior;
    real<lower=0> scale_sigma_prior;
    // Prior for beta (Normal)
    real loc_beta_prior;
    real<lower=0> scale_beta_prior;
    // Prior for sigma_beta (Normal)
    real loc_sigma_beta_prior;
    real<lower=0> scale_sigma_beta_prior;
}
transformed data{
    vector[2] xy[N];
    vector[2] zeros;
    for (i in 1:N){
        xy[i][1] = x[i];
        xy[i][2] = y[i];
    }

    zeros[1] = 0;
    zeros[2] = 0;
}
parameters {
    real beta;
    real<lower=0> sigma_beta;
    vector<lower=0>[S] sigma_y;
    vector<lower=0>[S] sigma_x;
    vector<lower=0>[S] tau;
    vector[S] beta_zc;
}
transformed parameters{
    cov_matrix[2] sigma_tmp[S];
    vector[S] beta_s;

    beta_s = beta + sigma_beta*beta_zc;
    for (s in 1:S){
        sigma_tmp[s][1,1] = square(sigma_x[s]) + square(tau[s]);
```

```
        sigma_tmp[s][1,2] = beta_s[s] * square(tau[s]);
        sigma_tmp[s][2,1] = beta_s[s] * square(tau[s]);
        sigma_tmp[s][2,2] = square(beta_s[s]) * square(tau[s]) + square(sigma_y[s]);
    }
}
model{
    beta ~ normal(loc_beta_prior, scale_beta_prior);
    sigma_beta ~ cauchy(loc_sigma_beta_prior, scale_sigma_beta_prior);
    beta_zc ~ normal(0, 1);
    sigma_y ~ cauchy(loc_sigma_prior, scale_sigma_prior);
    sigma_x ~ cauchy(loc_sigma_prior, scale_sigma_prior);
    tau ~ cauchy(loc_sigma_prior, scale_sigma_prior);

    for(s in 1:S){
        xy[(n_dat[s] + 1):n_dat[s+1]] ~ multi_normal(zeros, sigma_tmp[s]);
    }
}
generated quantities {
    real beta_pred;
    vector[S] corr;

    beta_pred = normal_rng(beta, sigma_beta);
    for(s in 1:S){
        corr[s]= sigma_tmp[s][1,2]/sqrt(sigma_tmp[s][1,1]*sigma_tmp[s][2,2]);
    }
}
```
