## Supplementary 2 for "Comparing and Validating Automated Tools for Individualized Electric Field Simulations in the Human Head"

April 17, 2019

# P03

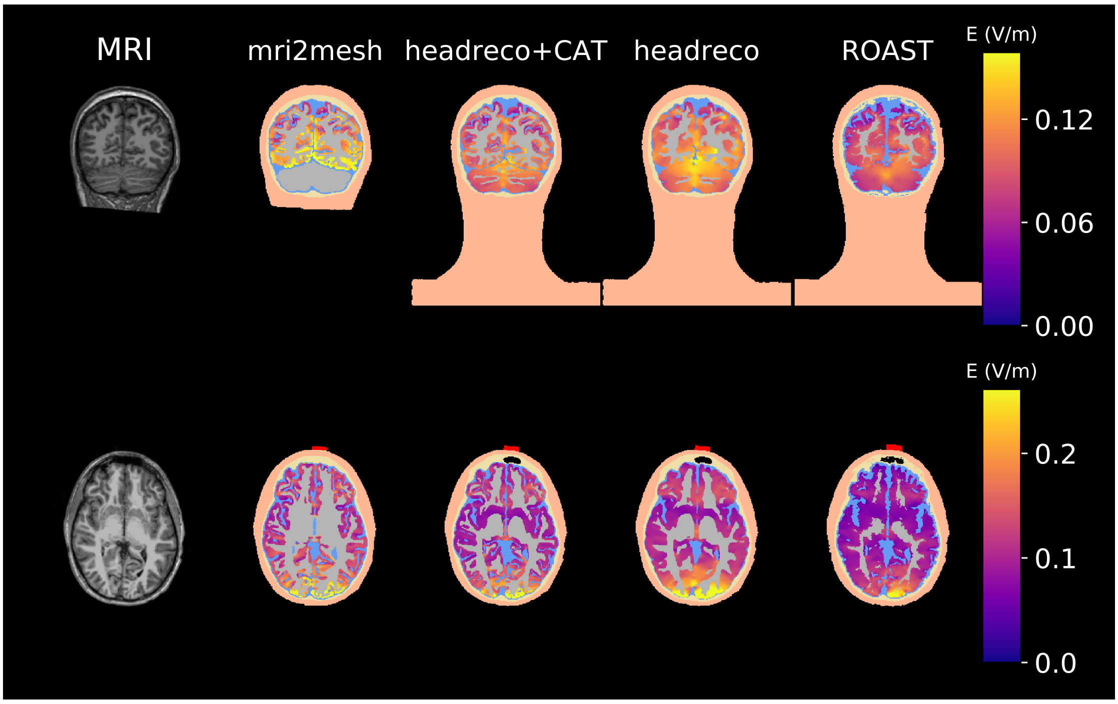


(a) T1-weighted image (first column), and segmentations with the electric field in gray matter

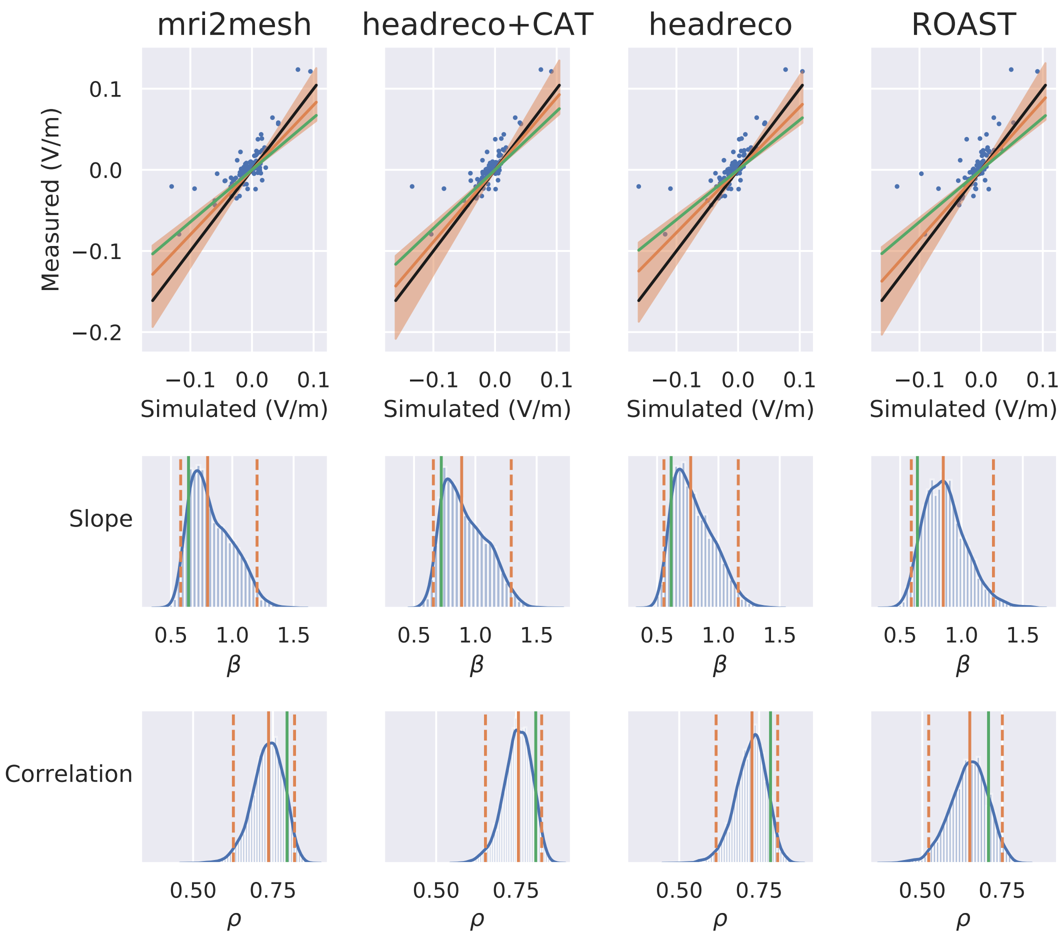


(b) Measured and simulated potential differences (first row), posterior probability for the slope $\beta$ (second row) and correlation $\rho$ (third row). Orange lines show the median and 95% compatibility interval obtained with the Bayesian errors-in-variables model. Green lines show the values obtained with standard regression analysis. The black line shows a slope of one.

Figure S.1: MRI, segmentations, simulations, recordings and fit for P03

# P04

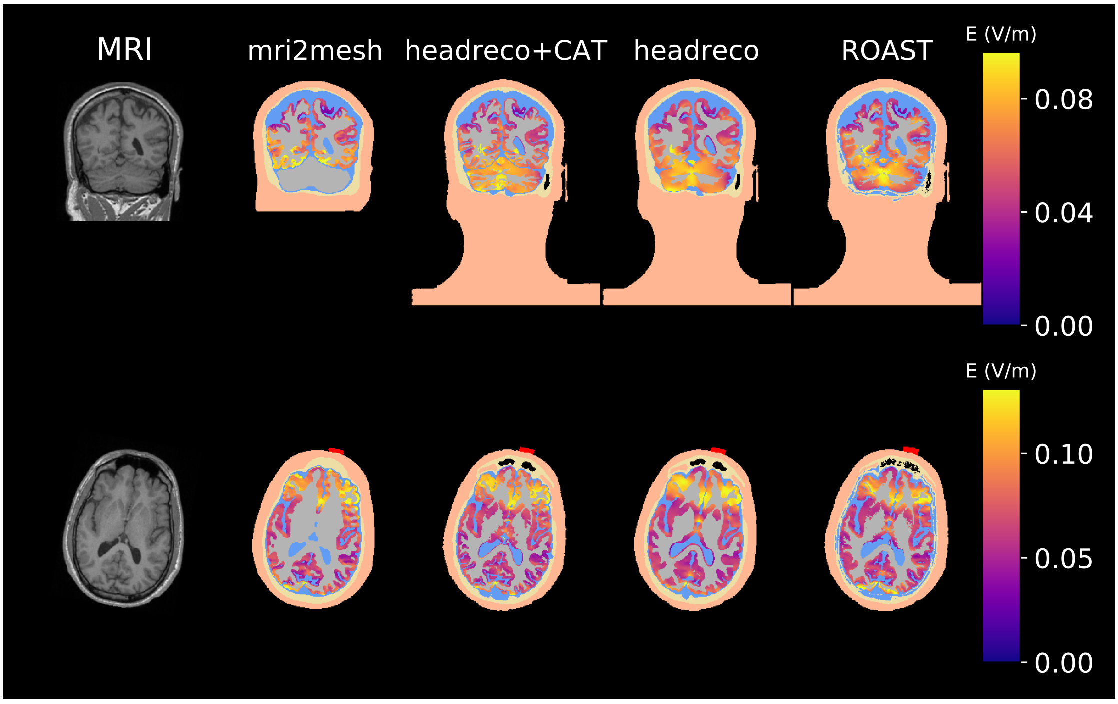


(a) T1-weighted image (first column), and segmentations with the electric field in gray matter

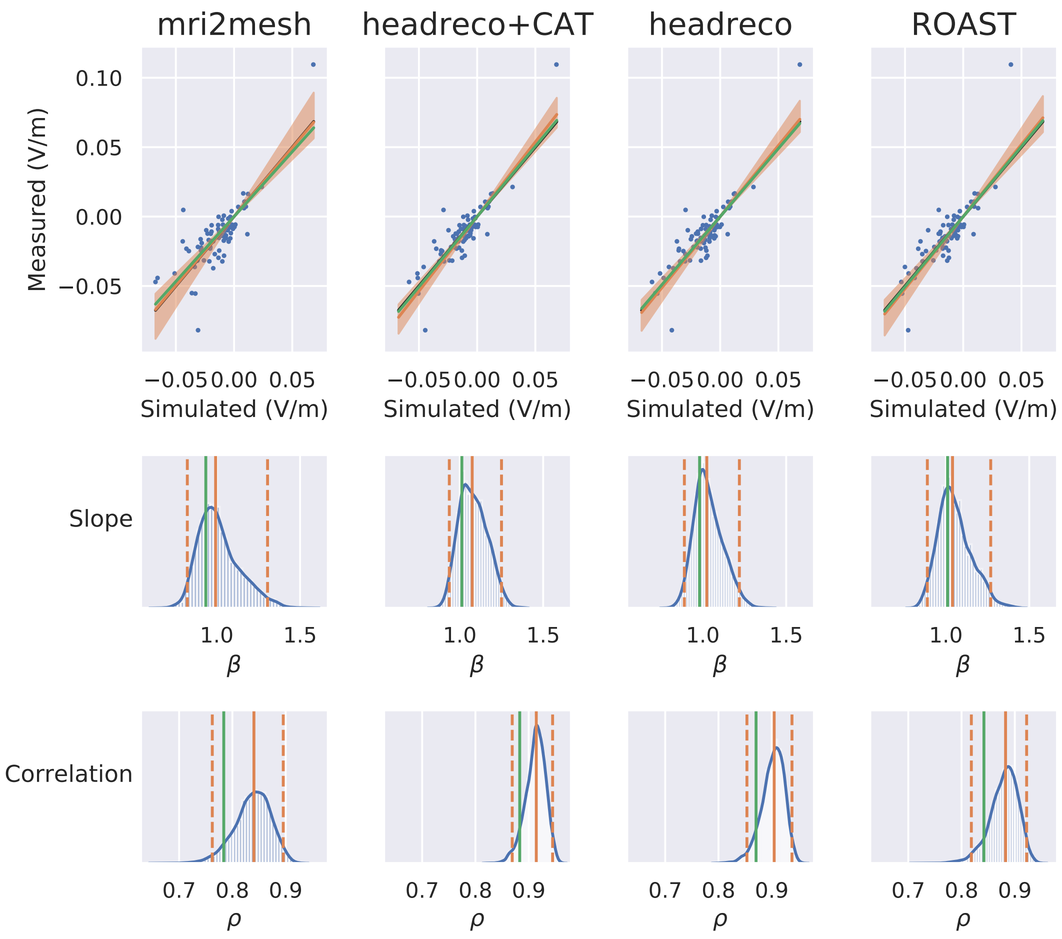


(b) Measured and simulated potential differences (first row), posterior probability for the slope $\beta$ (second row) and correlation $\rho$ (third row). Orange lines show the median and 95% compatibility interval obtained with the Bayesian errors-in-variables model. Green lines show the values obtained with standard regression analysis. The black line shows a slope of one.

Figure S.2: MRI, segmentations, simulations, recordings and fit for P04

# P05

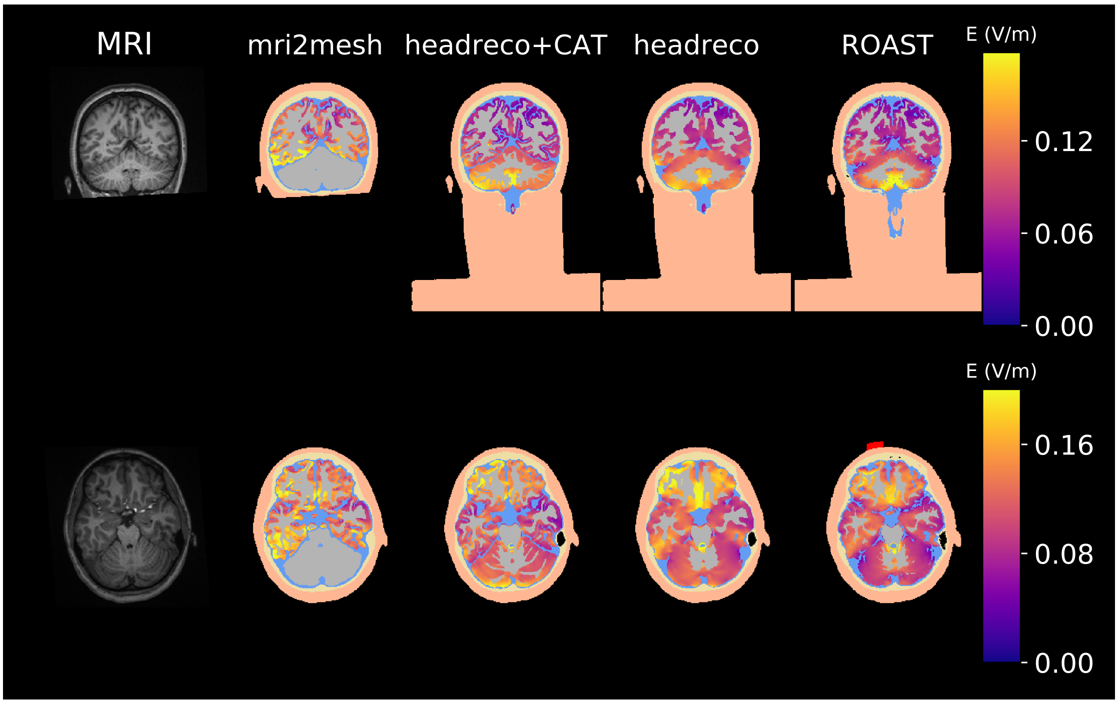


(a) T1-weighted image (first column), and segmentations with the electric field in gray matter

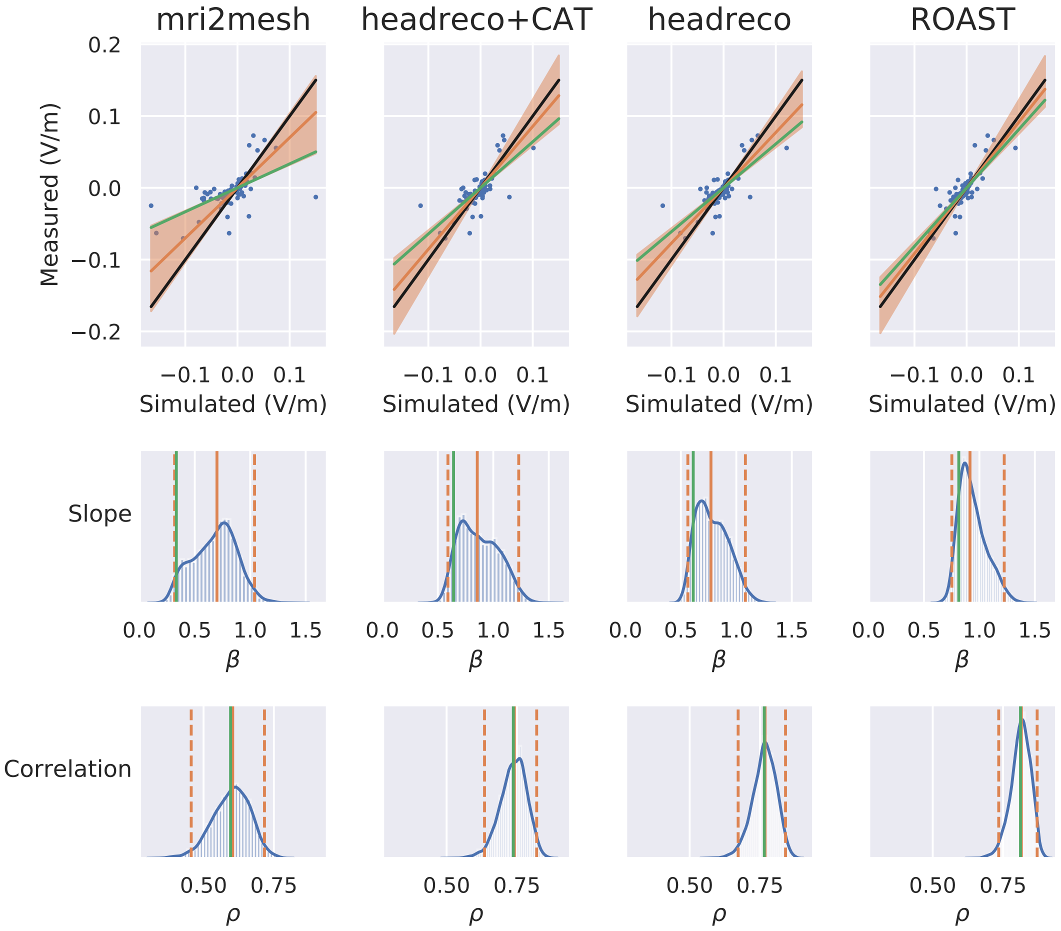


(b) Measured and simulated potential differences (first row), posterior probability for the slope $\beta$ (second row) and correlation $\rho$ (third row). Orange lines show the median and 95% compatibility interval obtained with the Bayesian errors-in-variables model. Green lines show the values obtained with standard regression analysis. The black line shows a slope of one.

Figure S.3: MRI, segmentations, simulations, recordings and fit for P05

# P06

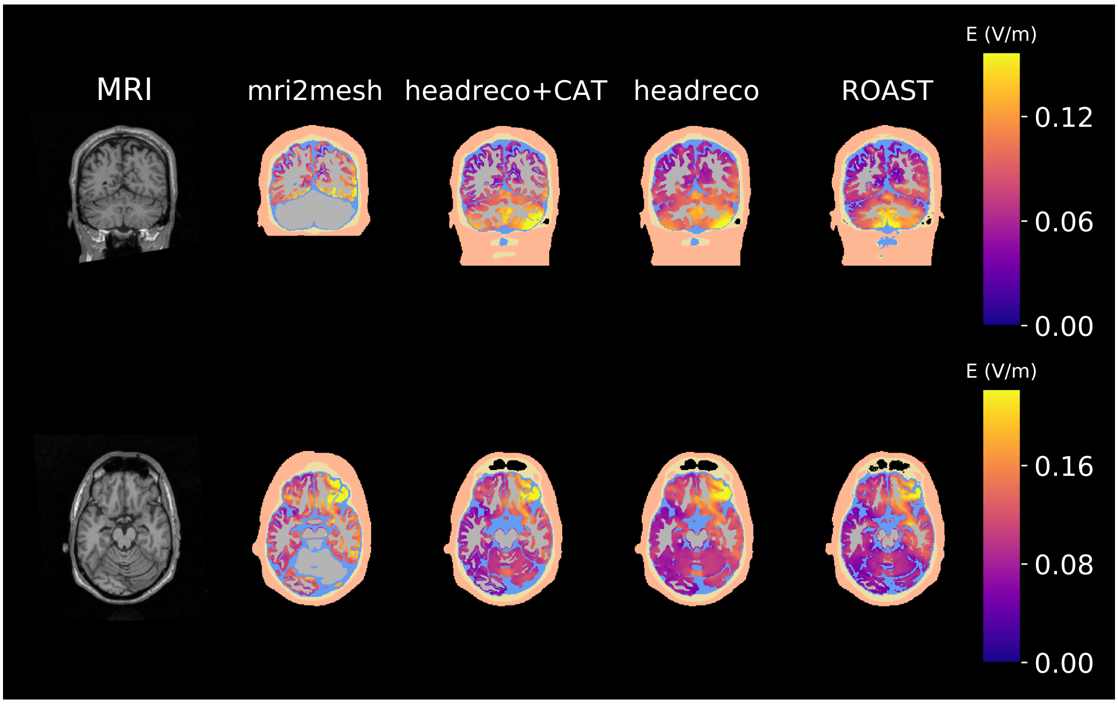


(a) T1-weighted image (first column), and segmentations with the electric field in gray matter

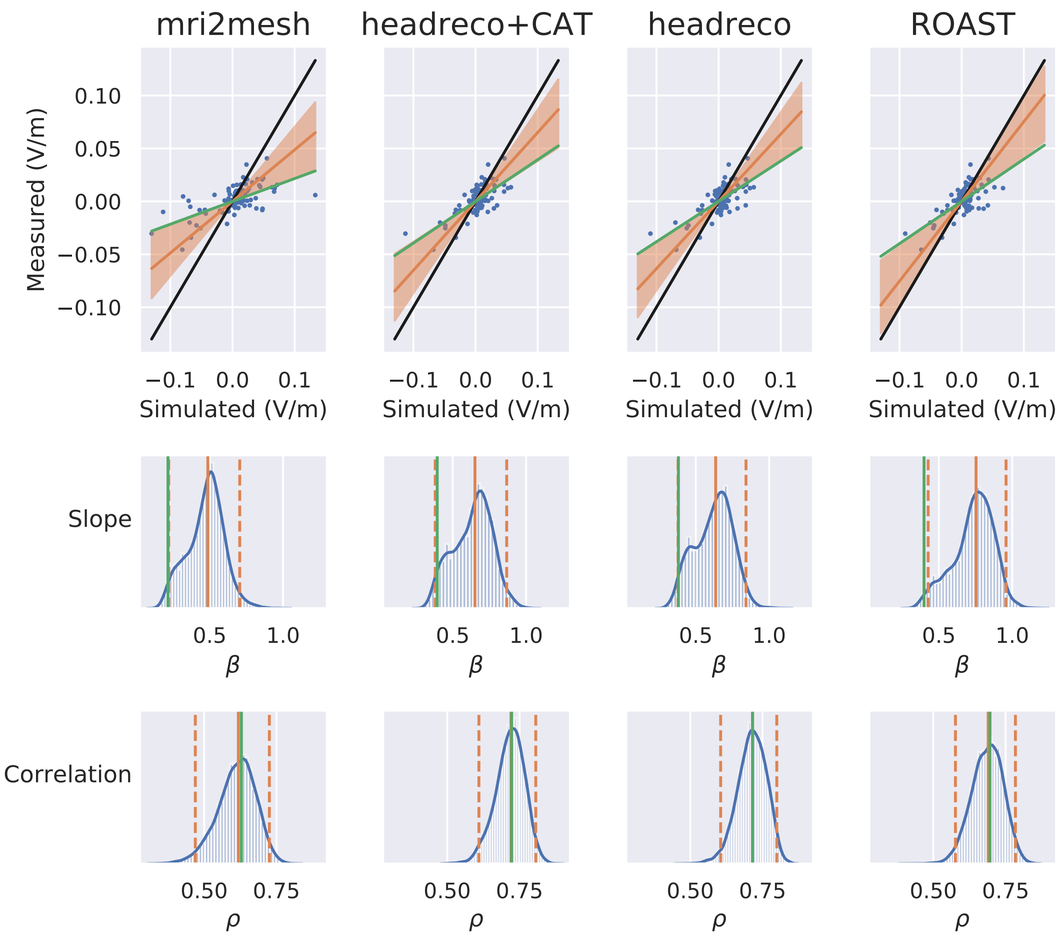


(b) Measured and simulated potential differences (first row), posterior probability for the slope $\beta$ (second row) and correlation $\rho$ (third row). Orange lines show the median and 95% compatibility interval obtained with the Bayesian errors-in-variables model. Green lines show the values obtained with standard regression analysis. The black line shows a slope of one.

Figure S.4: MRI, segmentations, simulations, recordings and fit for P06

# P07

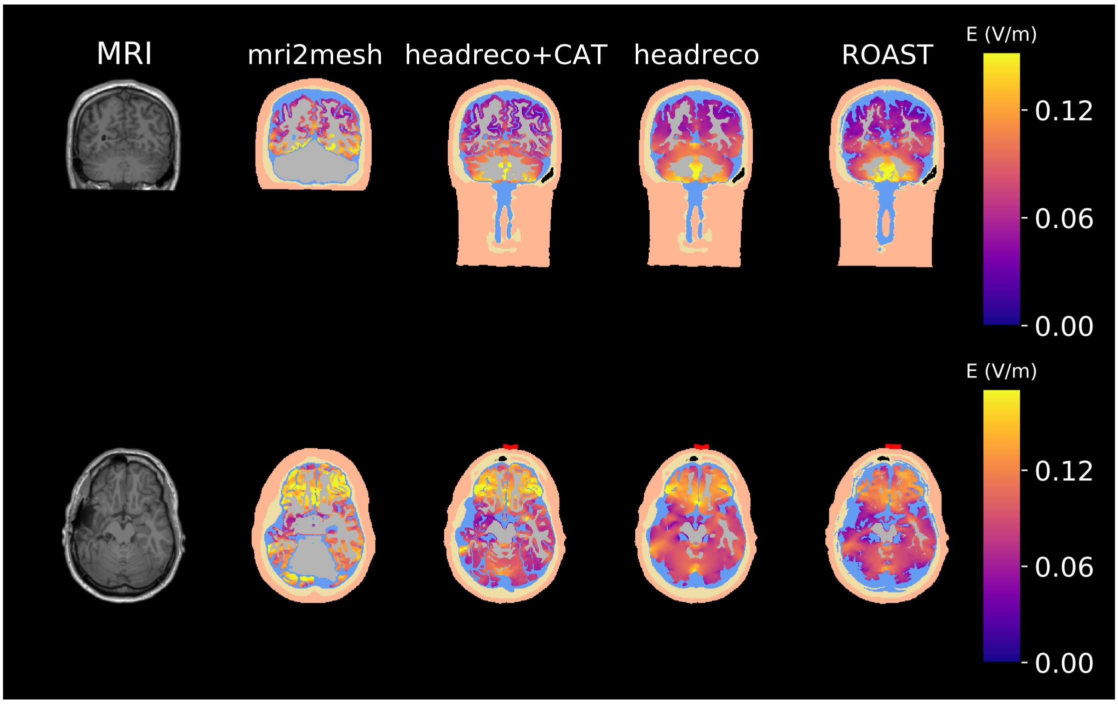


(a) T1-weighted image (first column), and segmentations with the electric field in gray matter

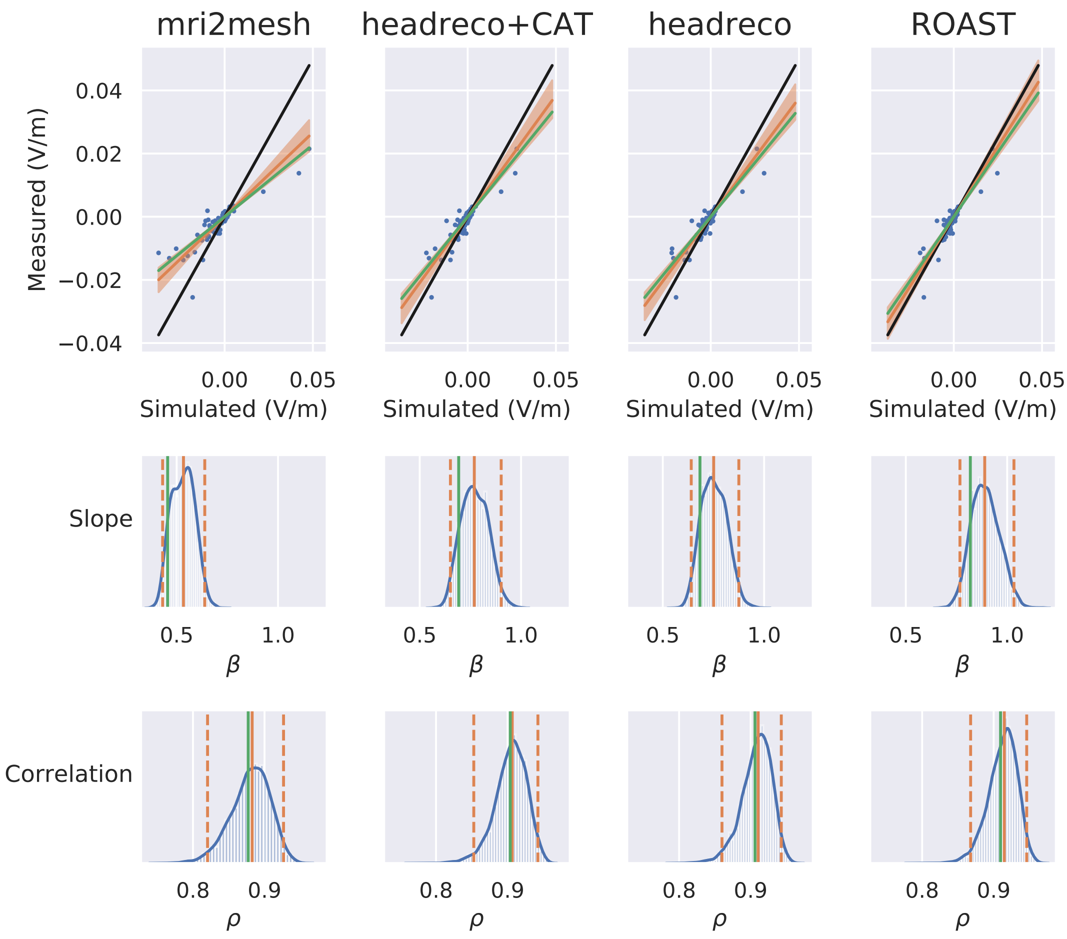


(b) Measured and simulated potential differences (first row), posterior probability for the slope $\beta$ (second row) and correlation $\rho$ (third row). Orange lines show the median and 95% compatibility interval obtained with the Bayesian errors-in-variables model. Green lines show the values obtained with standard regression analysis. The black line shows a slope of one.

Figure S.5: MRI, segmentations, simulations, recordings and fit for P07

# P08

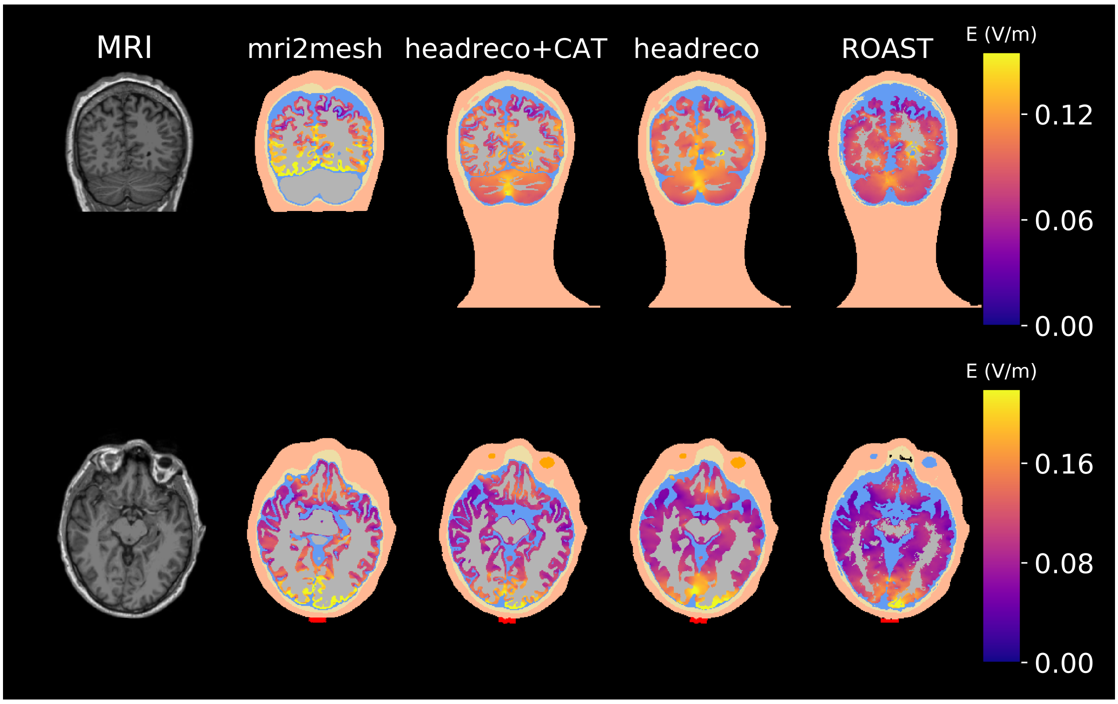


(a) T1-weighted image (first column), and segmentations with the electric field in gray matter

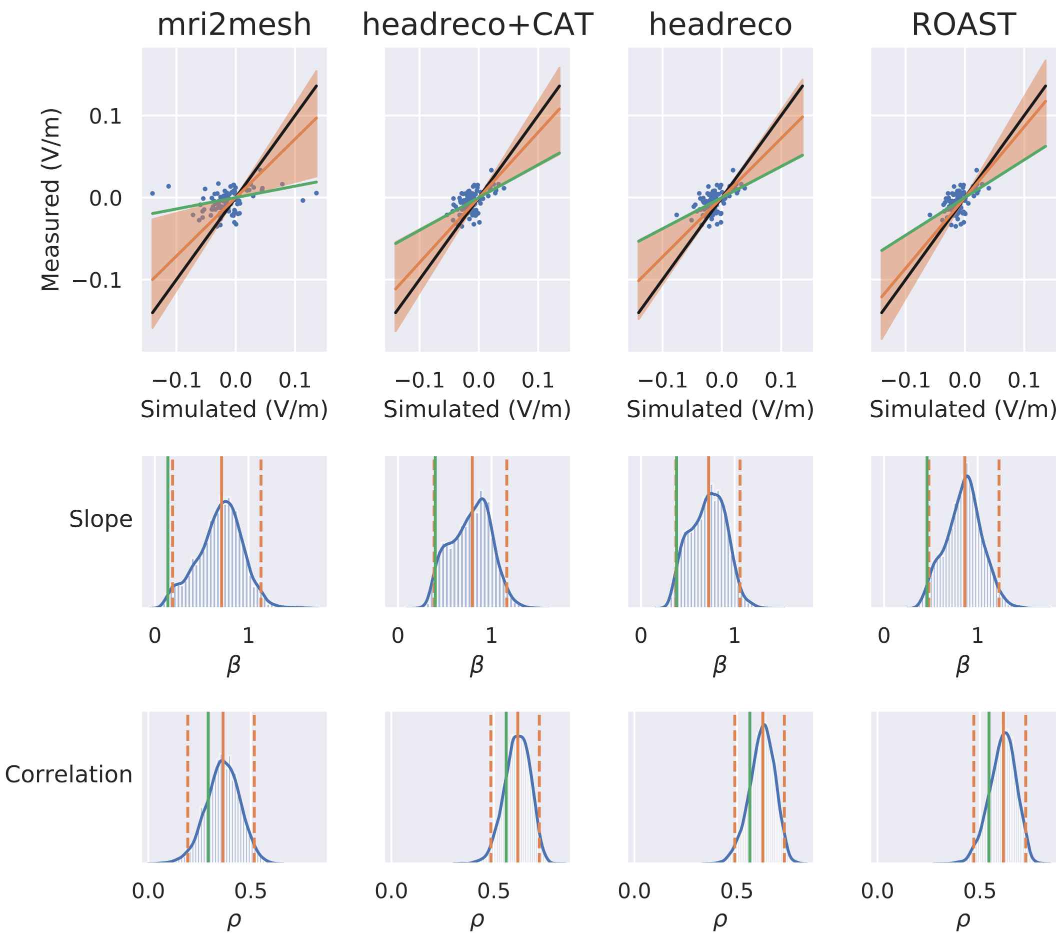


(b) Measured and simulated potential differences (first row), posterior probability for the slope $\beta$ (second row) and correlation $\rho$ (third row). Orange lines show the median and 95% compatibility interval obtained with the Bayesian errors-in-variables model. Green lines show the values obtained with standard regression analysis. The black line shows a slope of one.

Figure S.6: MRI, segmentations, simulations, recordings and fit for P08

# P09

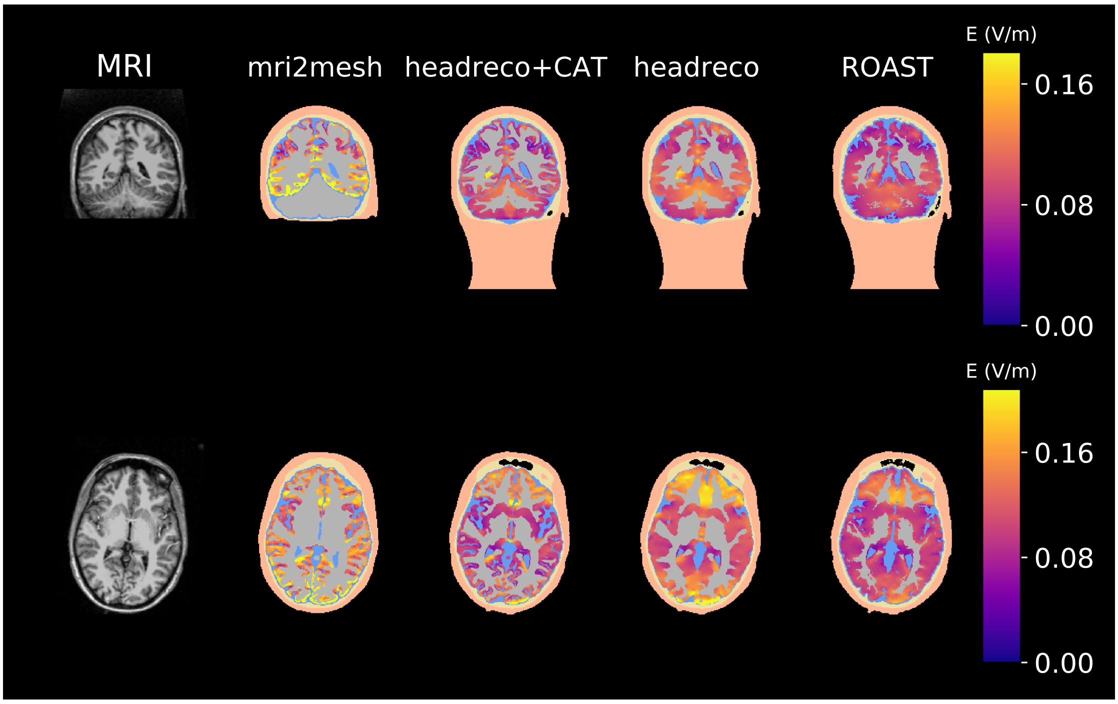


(a) T1-weighted image (first column), and segmentations with the electric field in gray matter

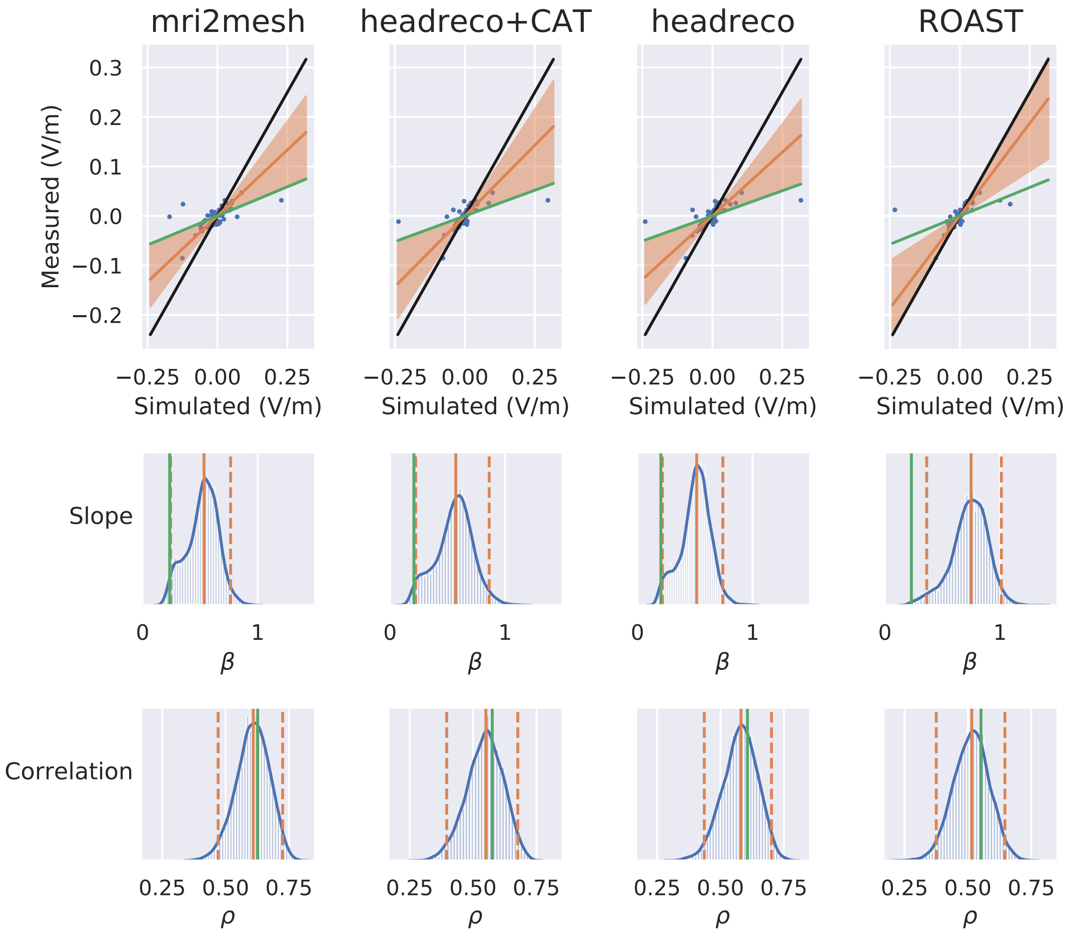


(b) Measured and simulated potential differences (first row), posterior probability for the slope $\beta$ (second row) and correlation $\rho$ (third row). Orange lines show the median and 95% compatibility interval obtained with the Bayesian errors-in-variables model. Green lines show the values obtained with standard regression analysis. The black line shows a slope of one.

Figure S.7: MRI, segmentations, simulations, recordings and fit for P09

# P010

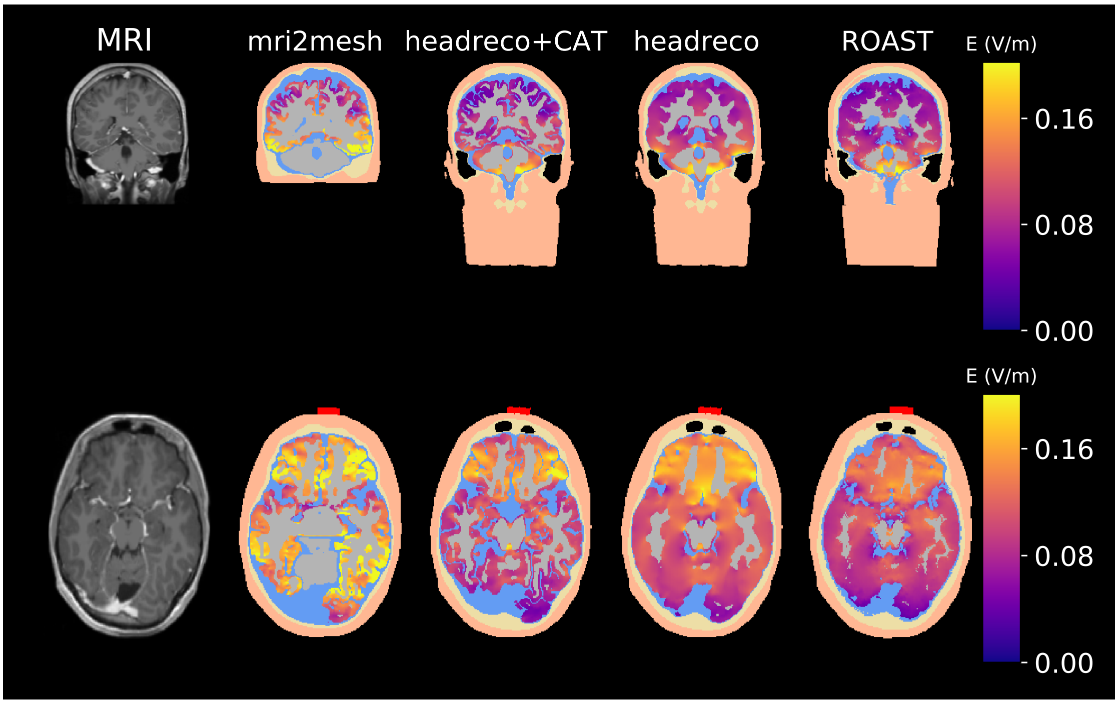


(a) T1-weighted image (first column), and segmentations with the electric field in gray matter

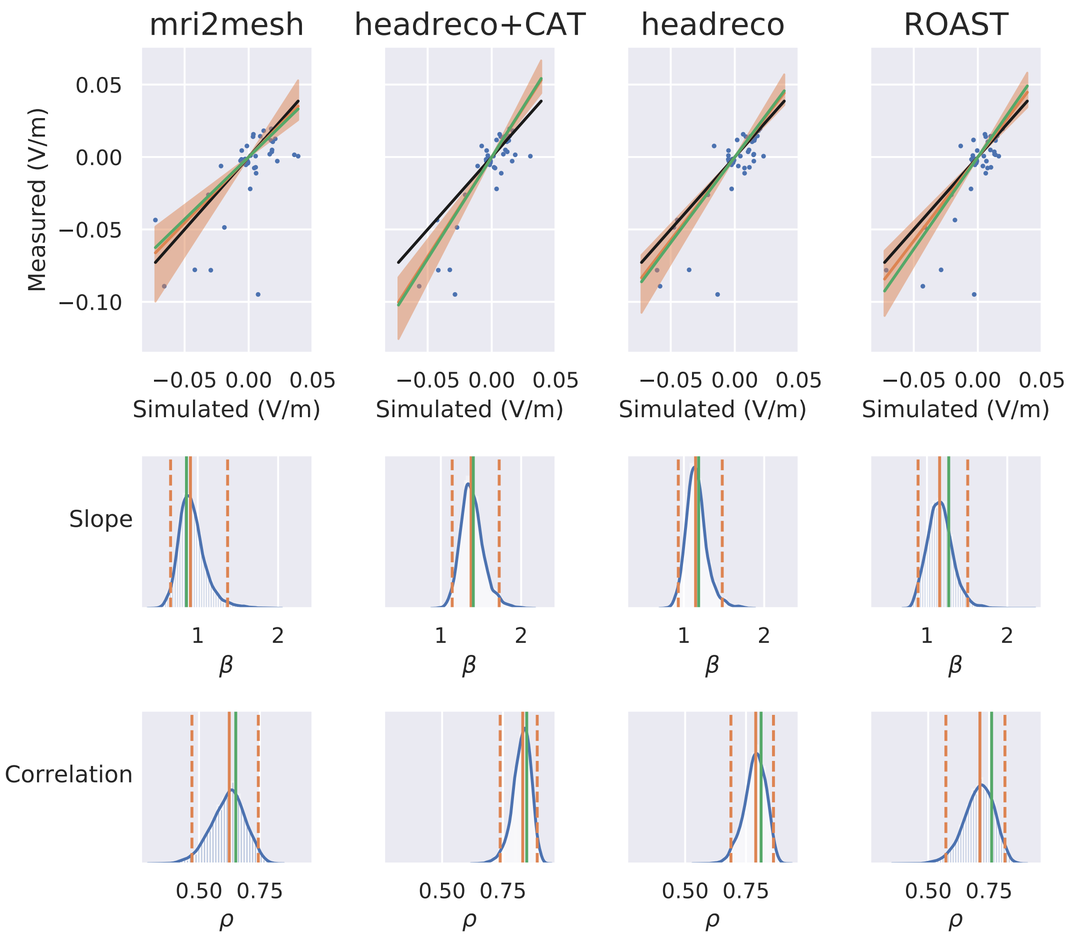


(b) Measured and simulated potential differences (first row), posterior probability for the slope $\beta$ (second row) and correlation $\rho$ (third row). Orange lines show the median and 95% compatibility interval obtained with the Bayesian errors-in-variables model. Green lines show the values obtained with standard regression analysis. The black line shows a slope of one.

Figure S.8: MRI, segmentations, simulations, recordings and fit for P010

# P011

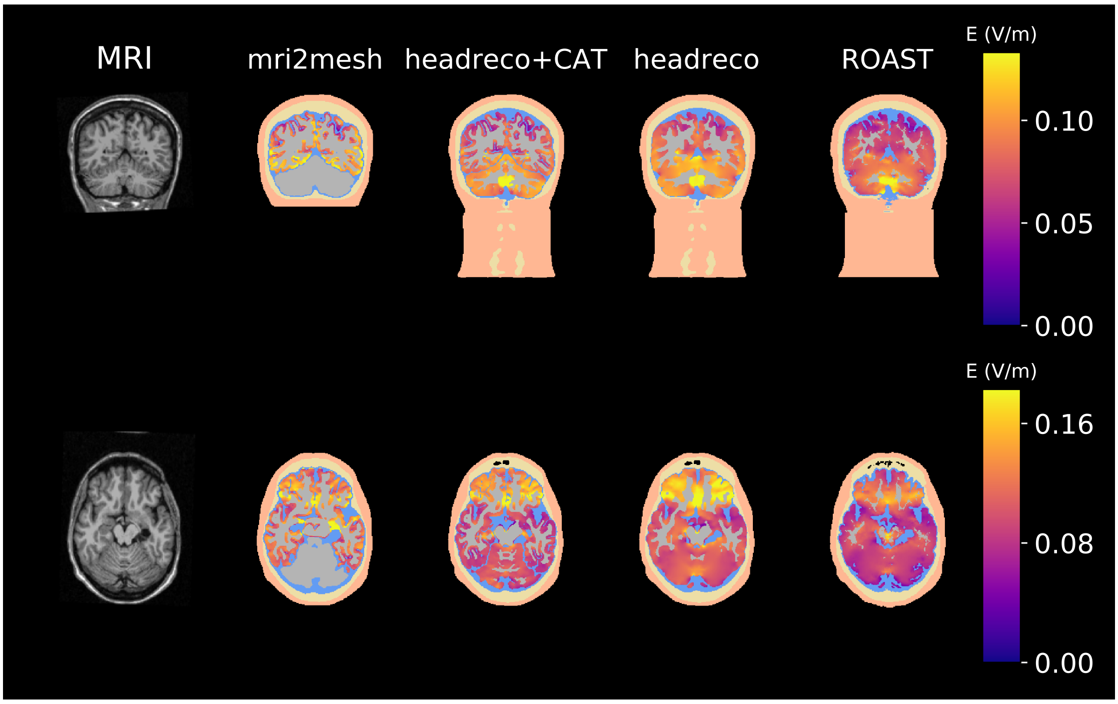


(a) T1-weighted image (first column), and segmentations with the electric field in gray matter

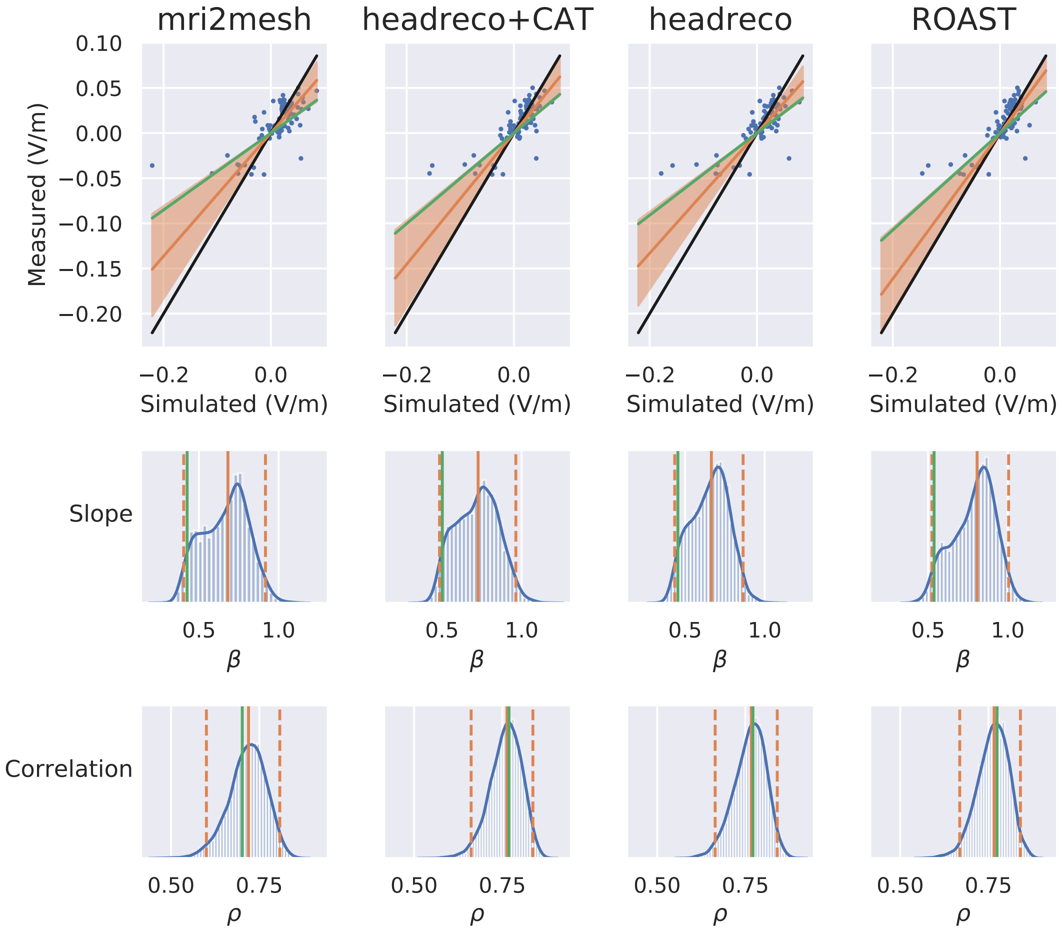


(b) Measured and simulated potential differences (first row), posterior probability for the slope $\beta$ (second row) and correlation $\rho$ (third row). Orange lines show the median and 95% compatibility interval obtained with the Bayesian errors-in-variables model. Green lines show the values obtained with standard regression analysis. The black line shows a slope of one.

Figure S.9: MRI, segmentations, simulations, recordings and fit for P011

# P013

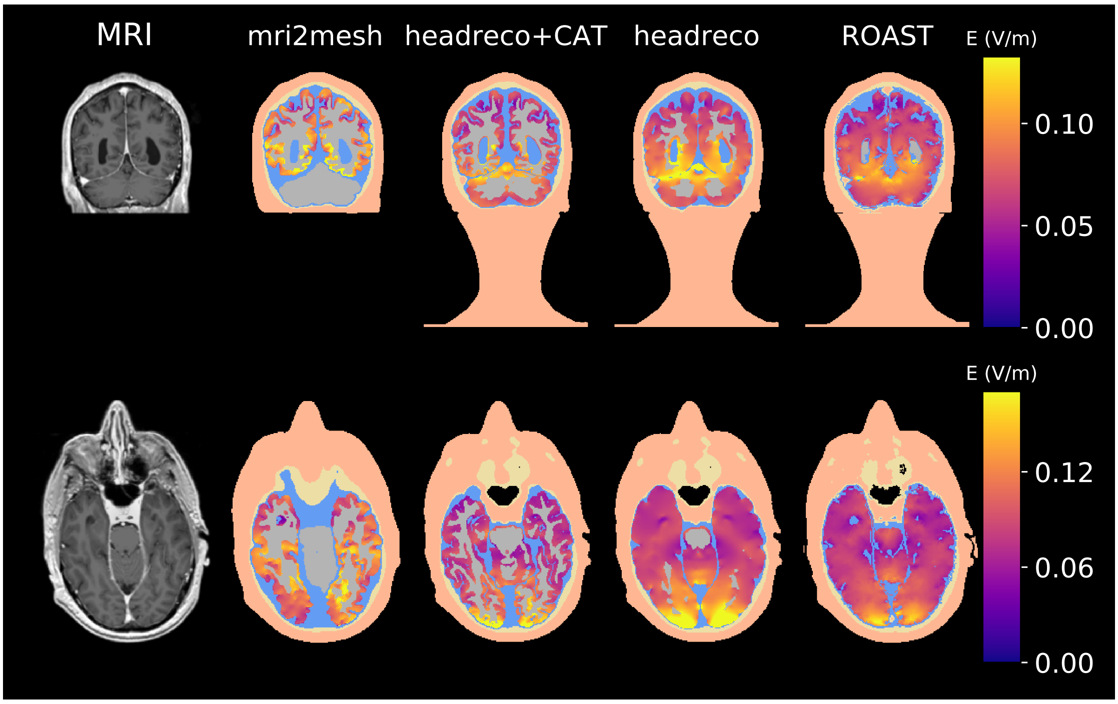


(a) T1-weighted image (first column), and segmentations with the electric field in gray matter

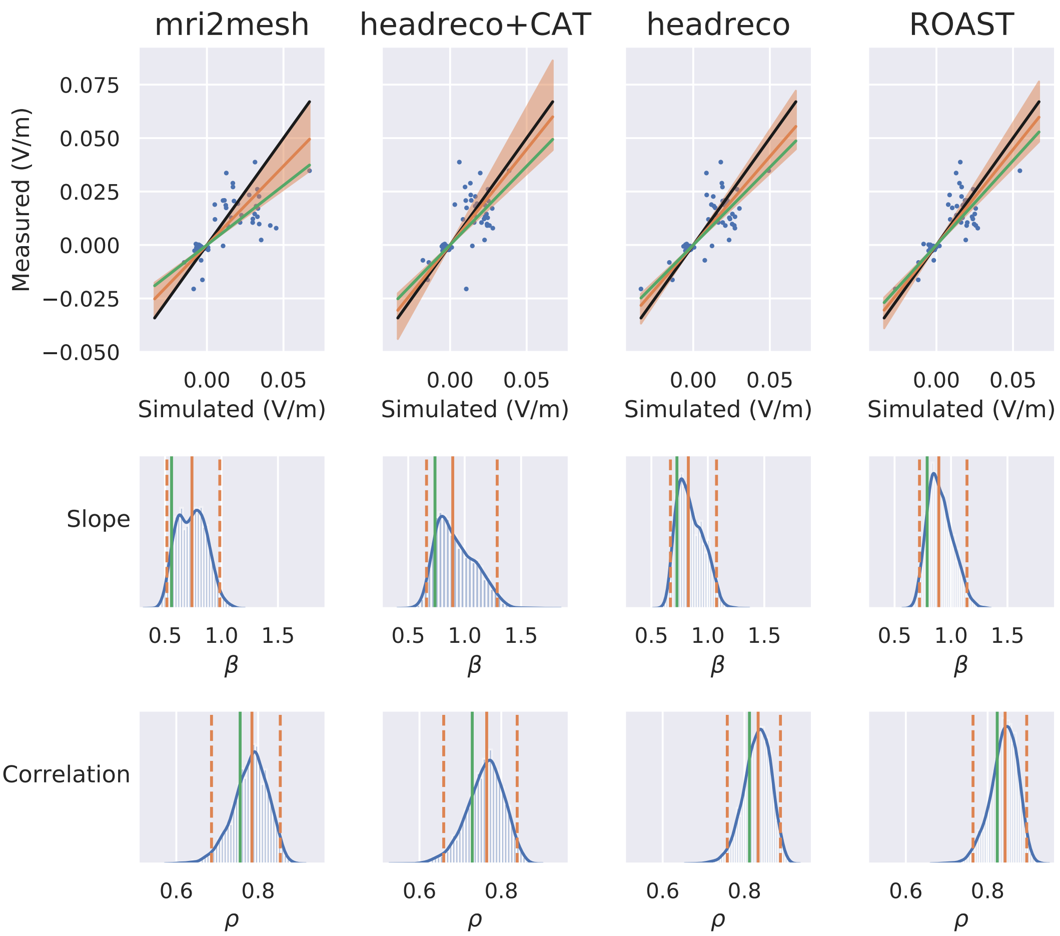


(b) Measured and simulated potential differences (first row), posterior probability for the slope $\beta$ (second row) and correlation $\rho$ (third row). Orange lines show the median and 95% compatibility interval obtained with the Bayesian errors-in-variables model. Green lines show the values obtained with standard regression analysis. The black line shows a slope of one.

Figure S.10: MRI, segmentations, simulations, recordings and fit for P013

# P014A

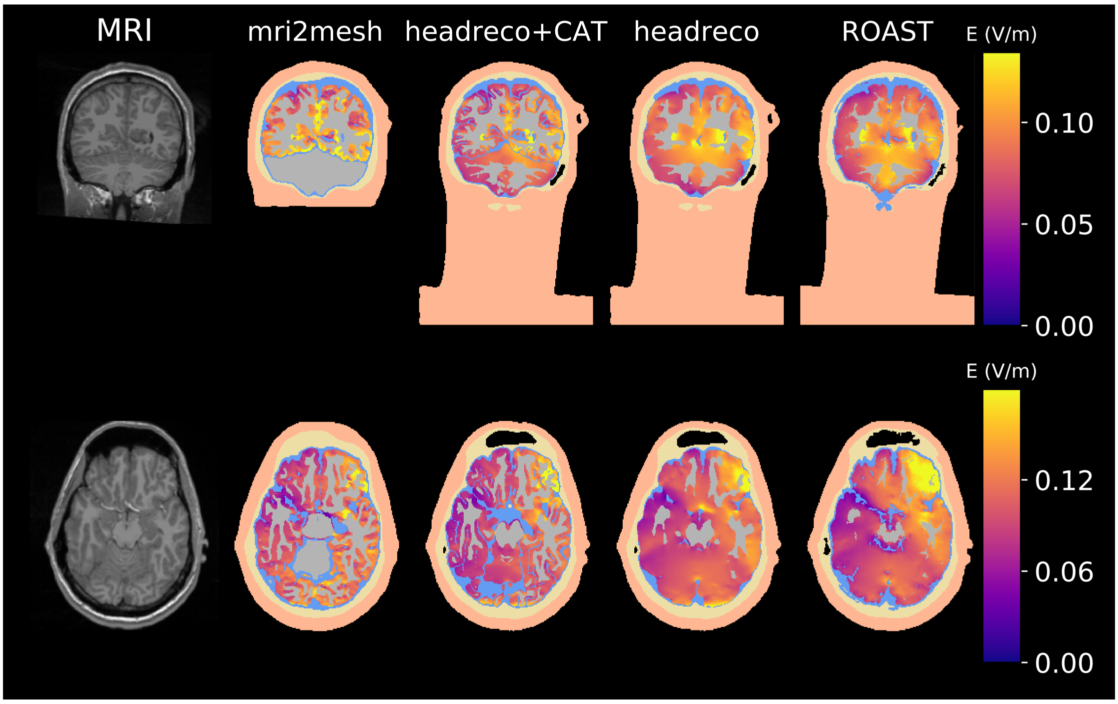


(a) T1-weighted image (first column), and segmentations with the electric field in gray matter

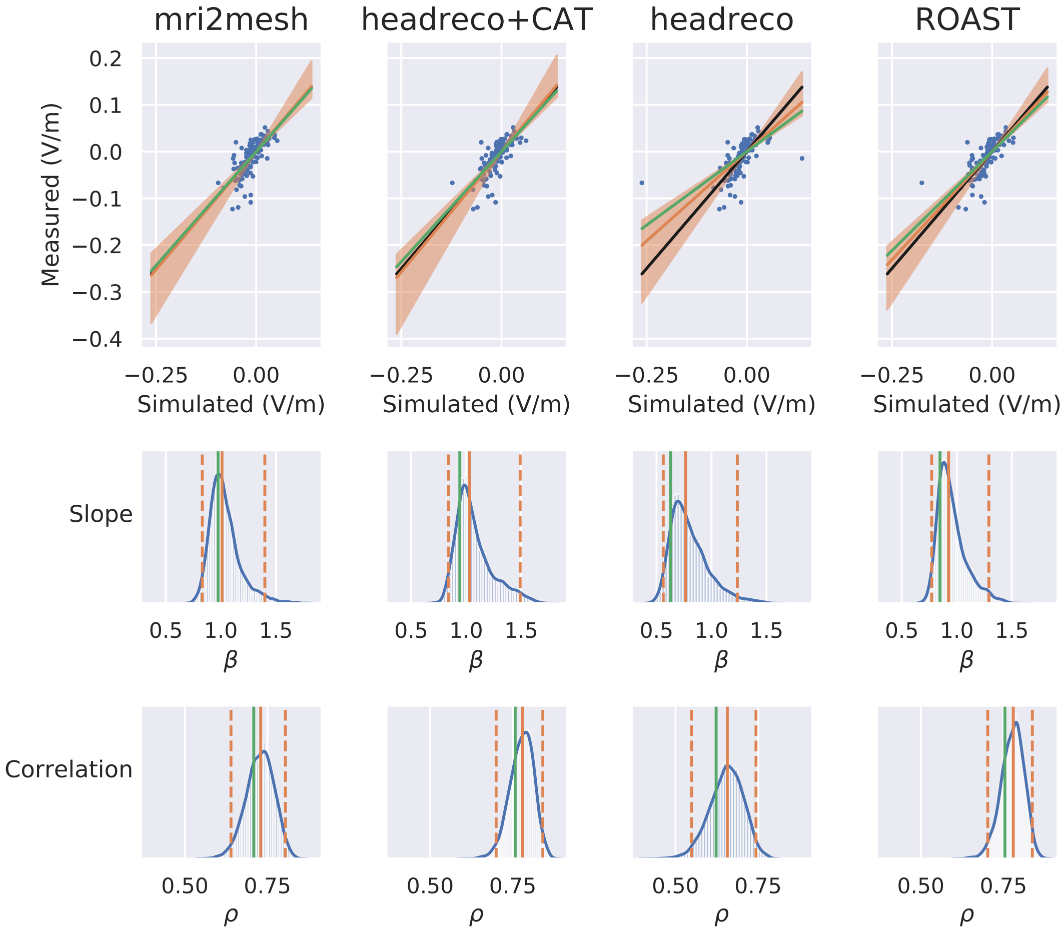


(b) Measured and simulated potential differences (first row), posterior probability for the slope $\beta$ (second row) and correlation $\rho$ (third row). Orange lines show the median and 95% compatibility interval obtained with the Bayesian errors-in-variables model. Green lines show the values obtained with standard regression analysis. The black line shows a slope of one.

Figure S.11: MRI, segmentations, simulations, recordings and fit for P014A

# P014B

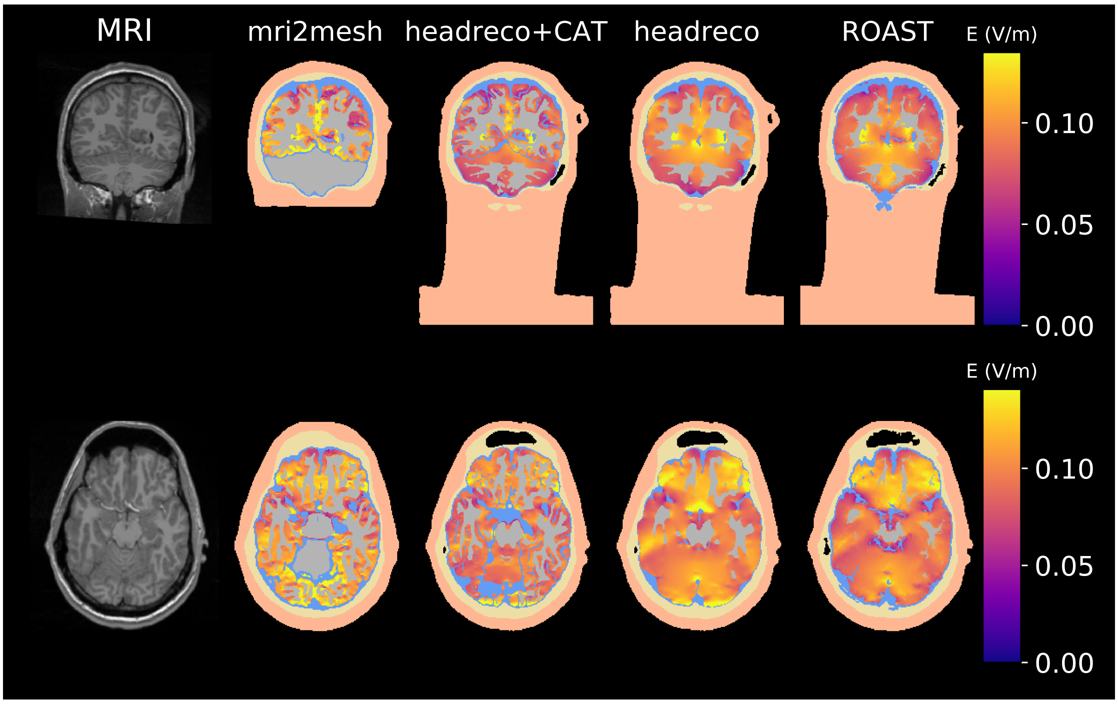


(a) T1-weighted image (first column), and segmentations with the electric field in gray matter

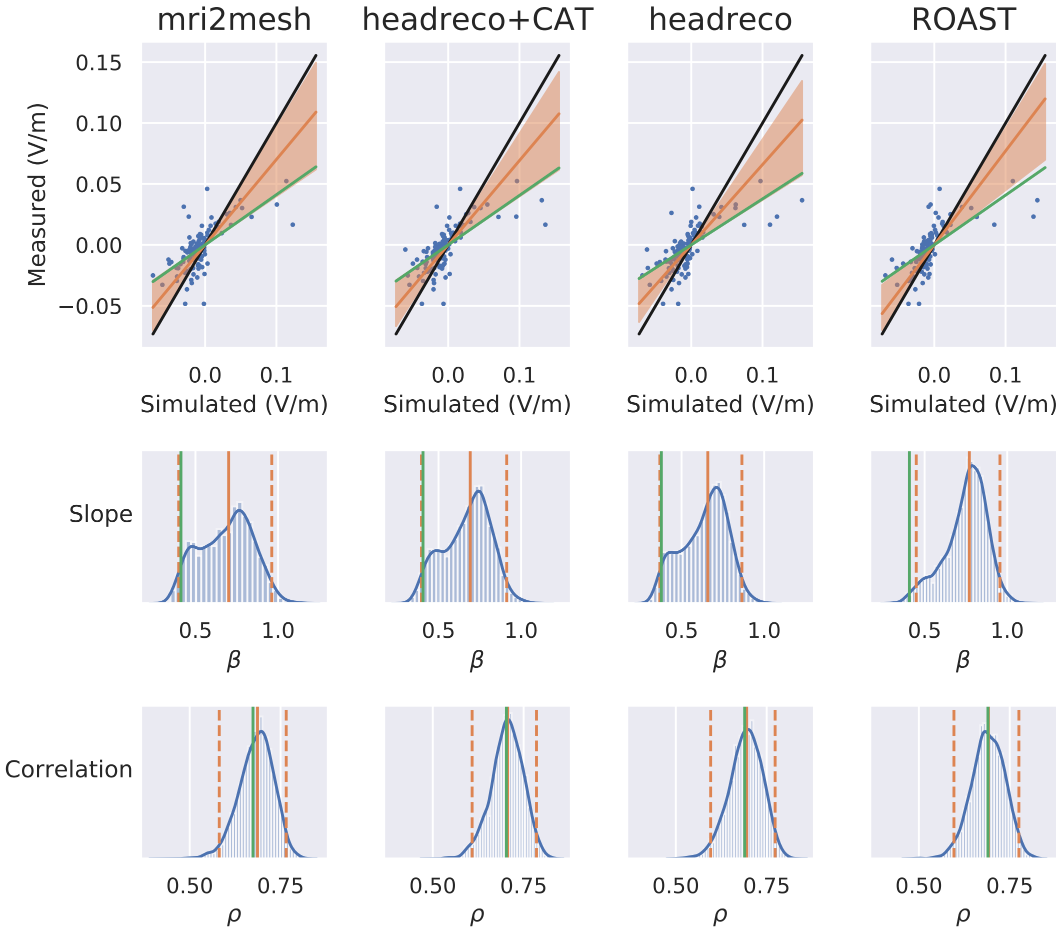


(b) Measured and simulated potential differences (first row), posterior probability for the slope $\beta$ (second row) and correlation $\rho$ (third row). Orange lines show the median and 95% compatibility interval obtained with the Bayesian errors-in-variables model. Green lines show the values obtained with standard regression analysis. The black line shows a slope of one.

Figure S.12: MRI, segmentations, simulations, recordings and fit for P014B

# P014C

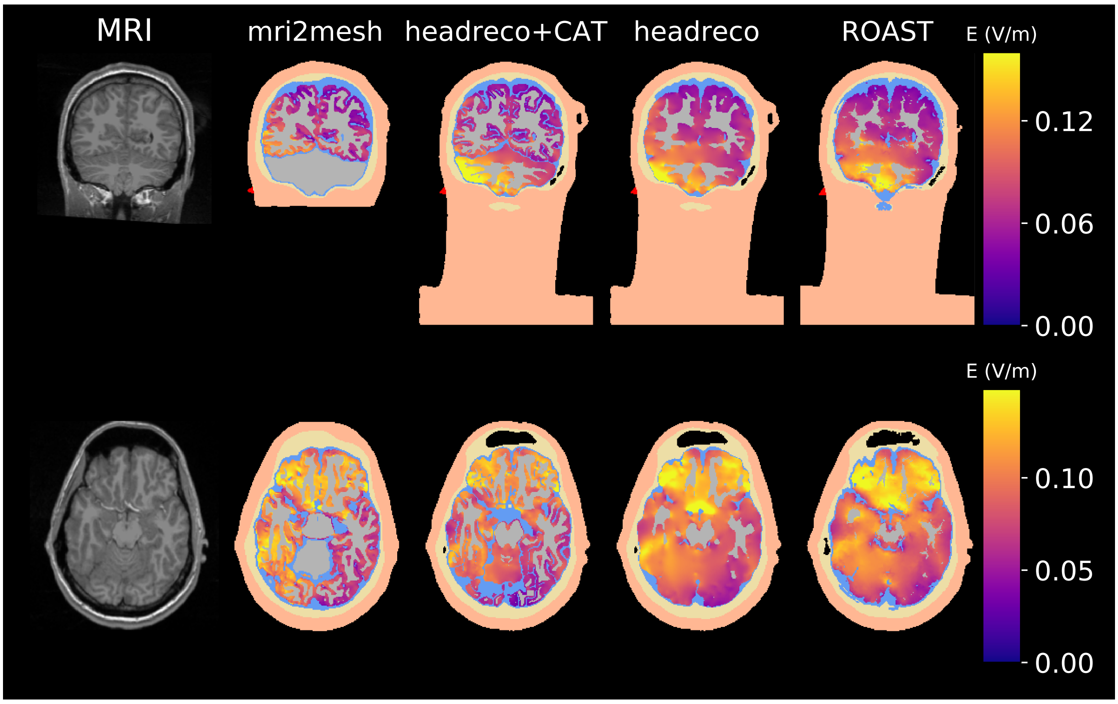


(a) T1-weighted image (first column), and segmentations with the electric field in gray matter

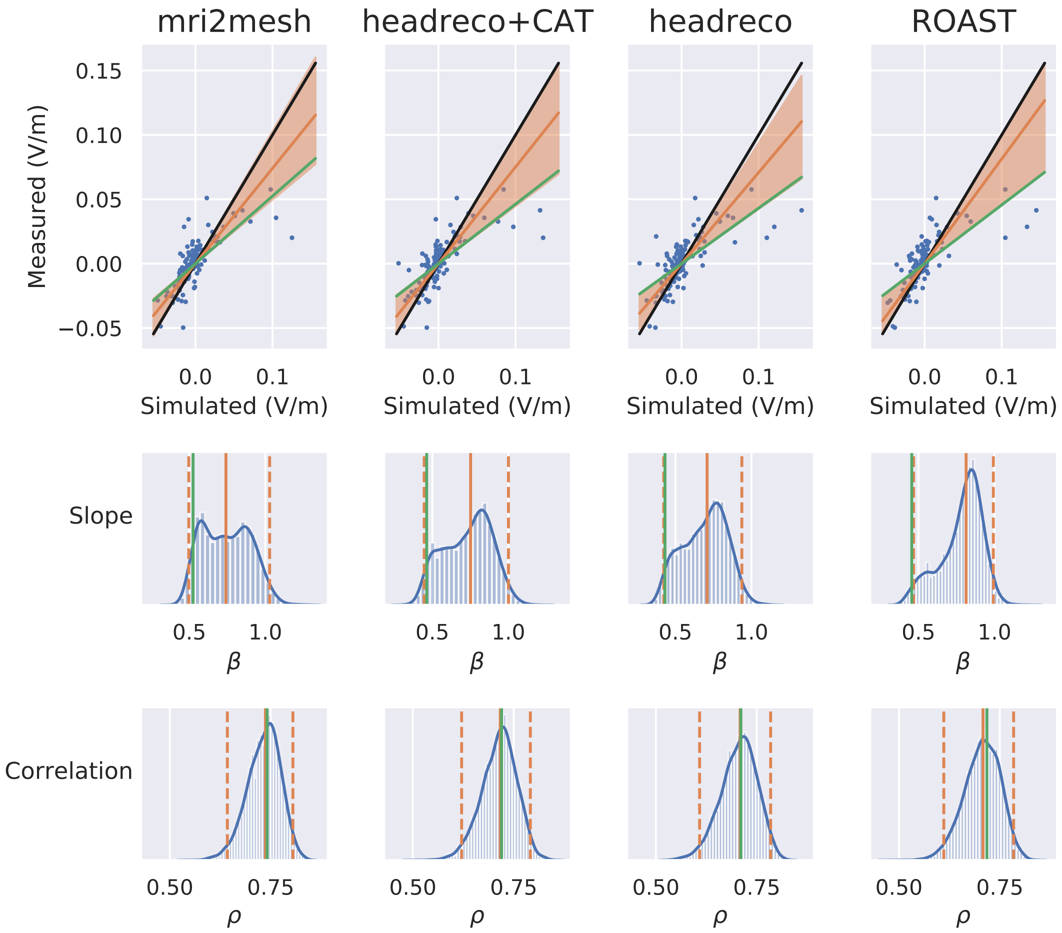


(b) Measured and simulated potential differences (first row), posterior probability for the slope $\beta$ (second row) and correlation $\rho$ (third row). Orange lines show the median and 95% compatibility interval obtained with the Bayesian errors-in-variables model. Green lines show the values obtained with standard regression analysis. The black line shows a slope of one.

Figure S.13: MRI, segmentations, simulations, recordings and fit for P014C

# P014D

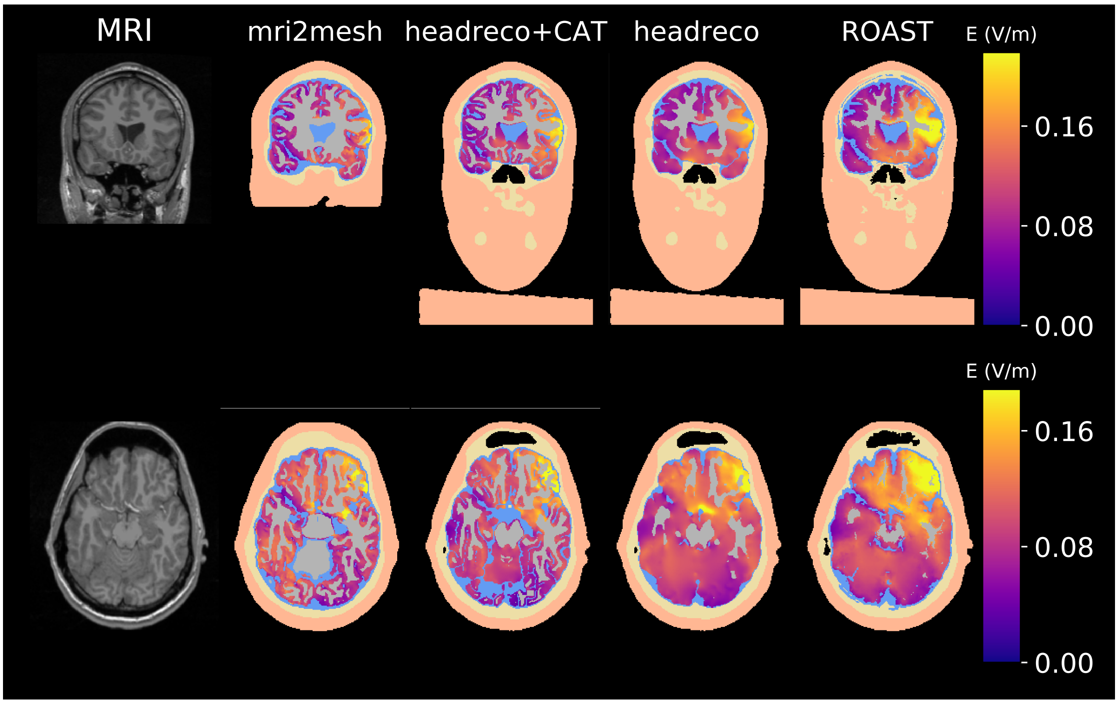


(a) T1-weighted image (first column), and segmentations with the electric field in gray matter

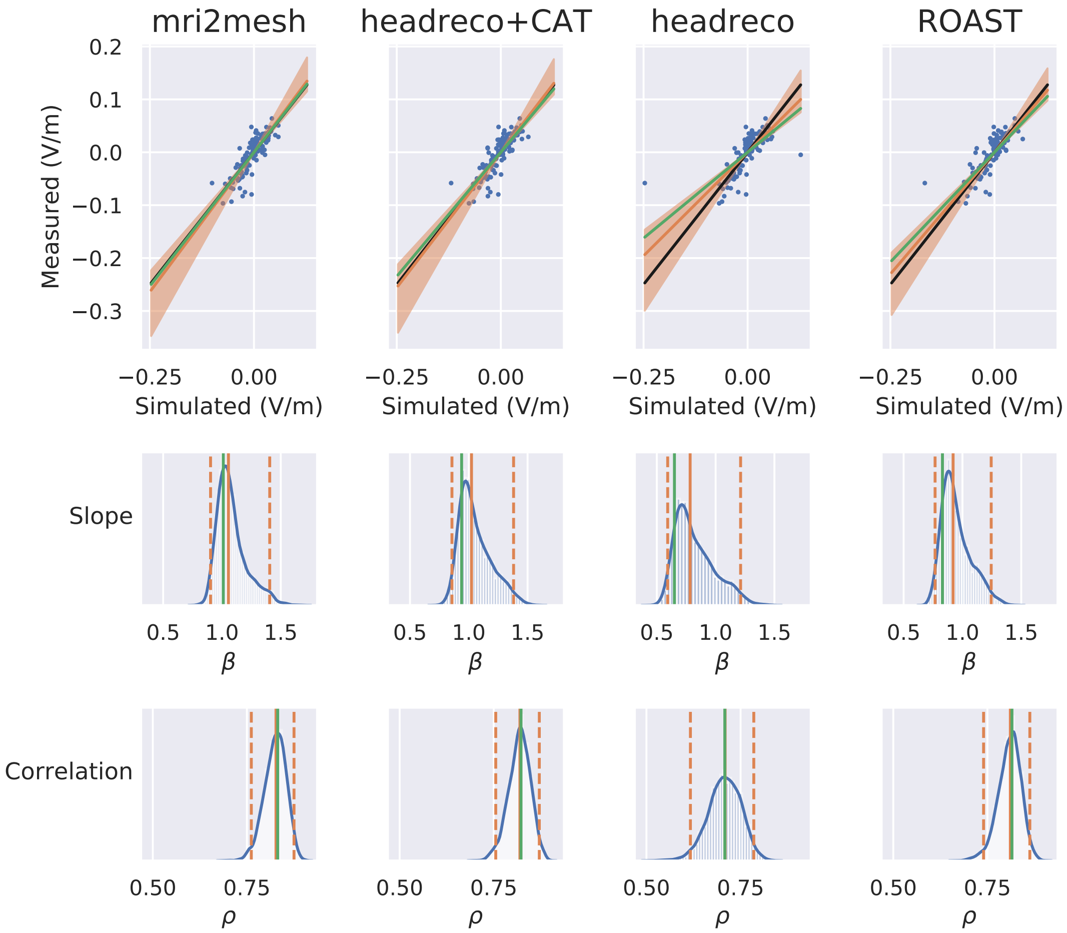


(b) Measured and simulated potential differences (first row), posterior probability for the slope $\beta$ (second row) and correlation $\rho$ (third row). Orange lines show the median and 95% compatibility interval obtained with the Bayesian errors-in-variables model. Green lines show the values obtained with standard regression analysis. The black line shows a slope of one.

Figure S.14: MRI, segmentations, simulations, recordings and fit for P014D

# P015

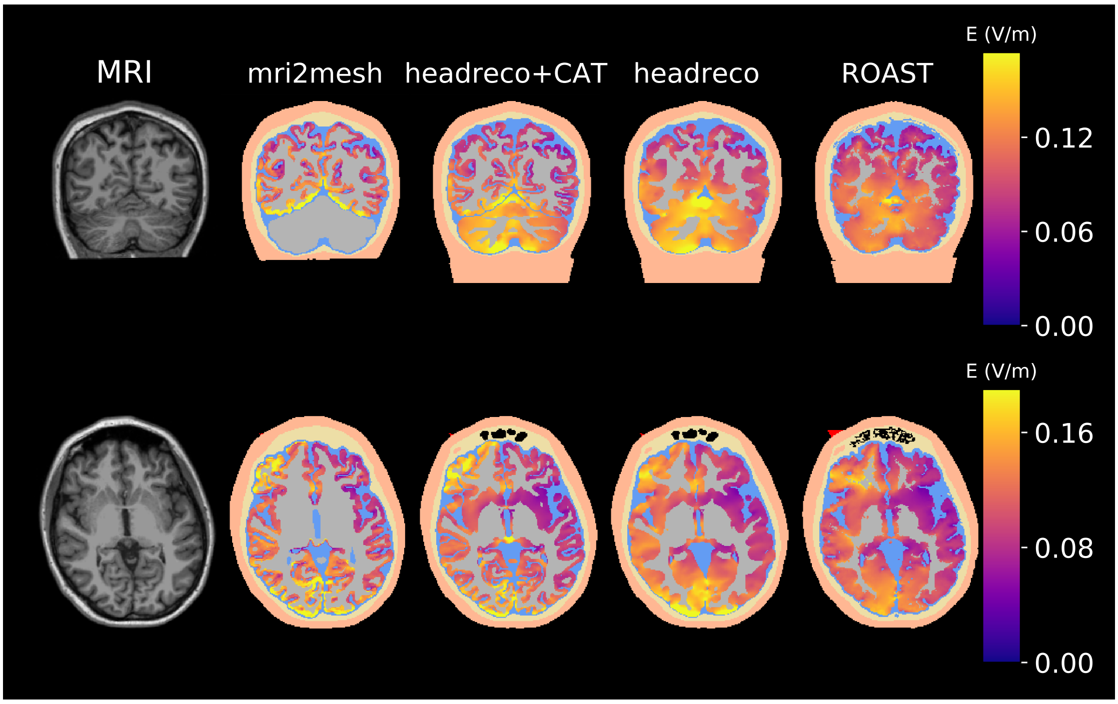


(a) T1-weighted image (first column), and segmentations with the electric field in gray matter

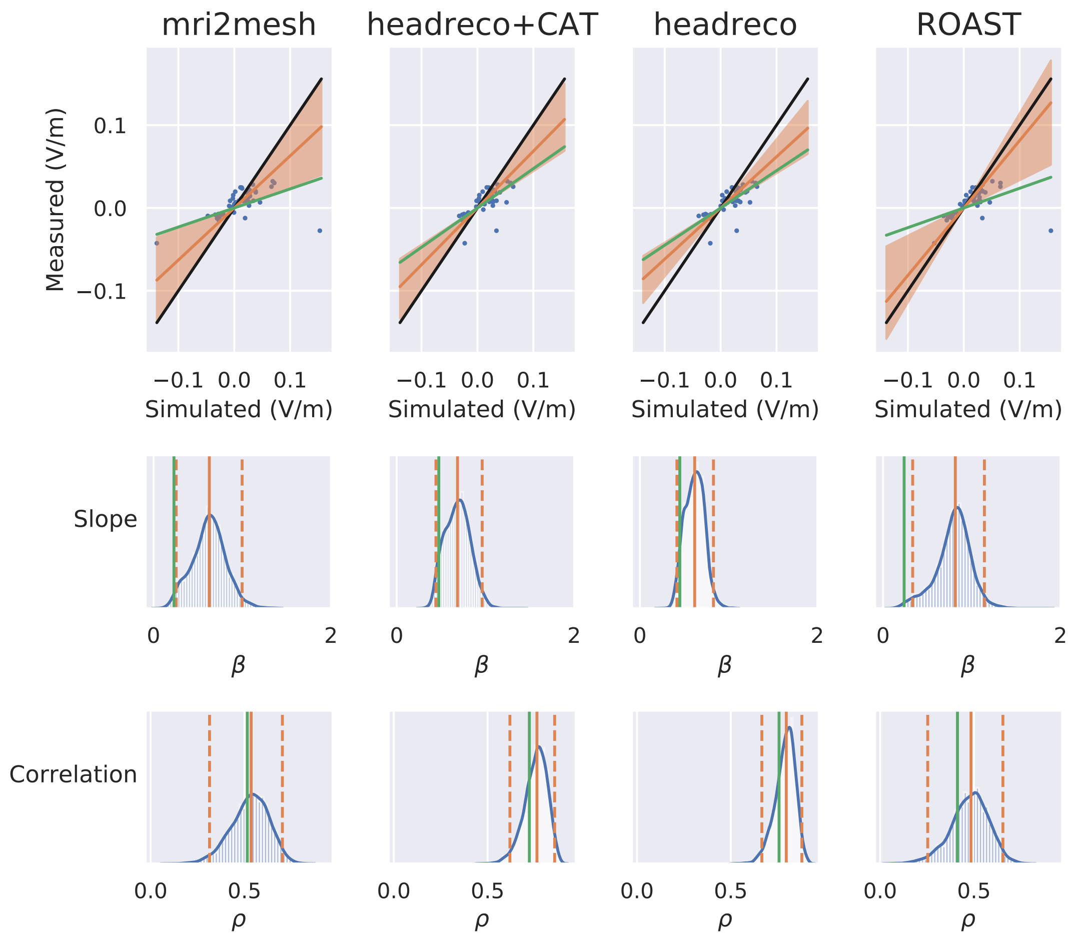


(b) Measured and simulated potential differences (first row), posterior probability for the slope $\beta$ (second row) and correlation $\rho$ (third row). Orange lines show the median and 95% compatibility interval obtained with the Bayesian errors-in-variables model. Green lines show the values obtained with standard regression analysis. The black line shows a slope of one.

Figure S.15: MRI, segmentations, simulations, recordings and fit for P015

# P016

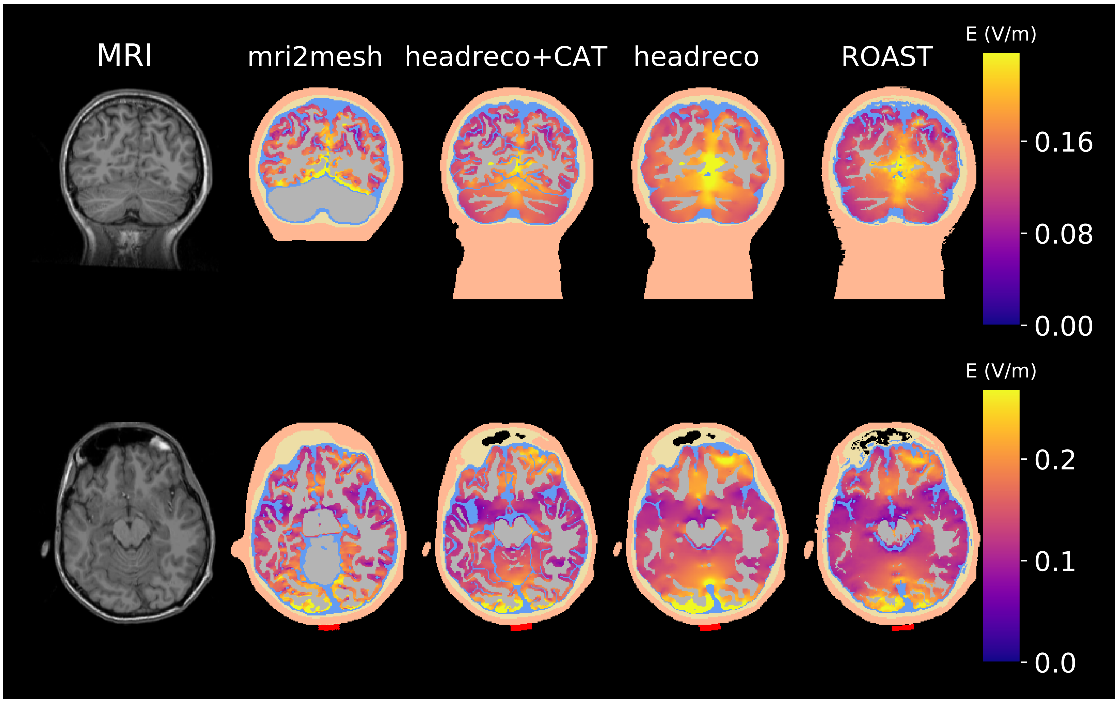


(a) T1-weighted image (first column), and segmentations with the electric field in gray matter

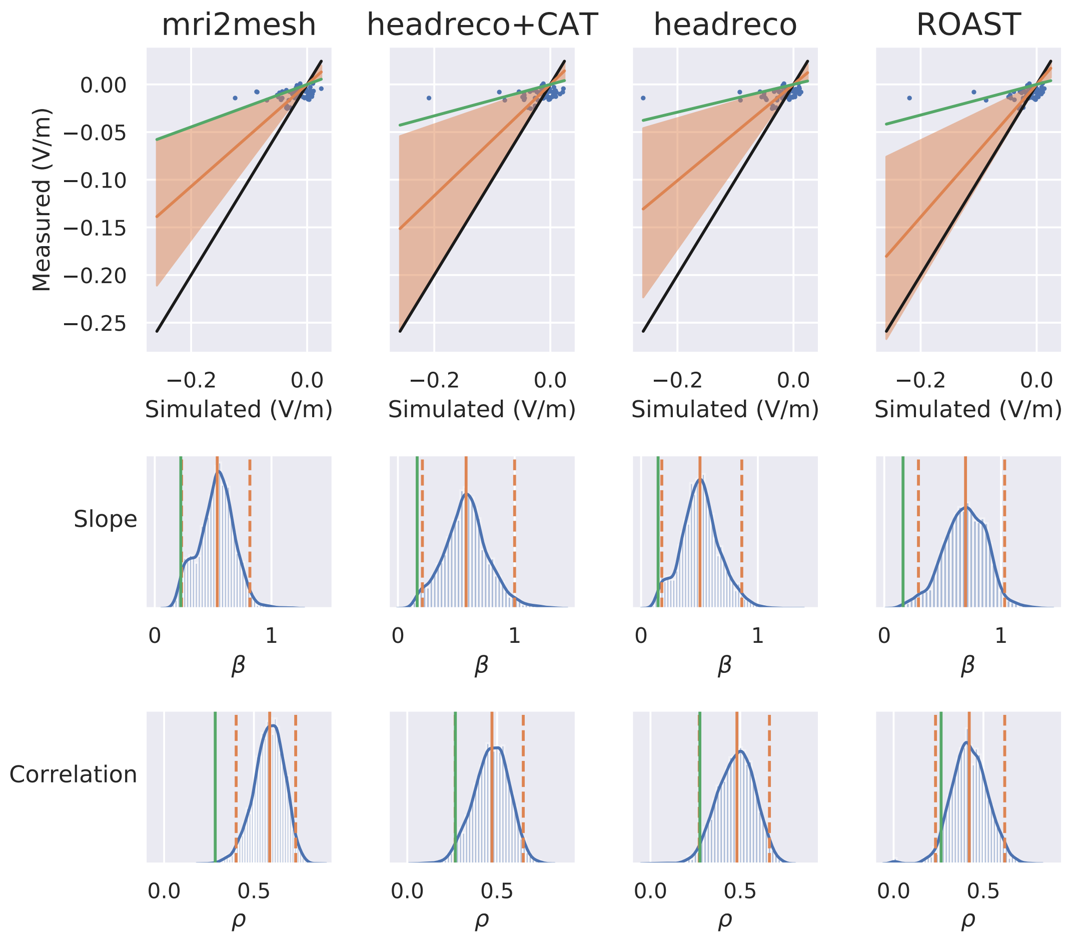


(b) Measured and simulated potential differences (first row), posterior probability for the slope $\beta$ (second row) and correlation $\rho$ (third row). Orange lines show the median and 95% compatibility interval obtained with the Bayesian errors-in-variables model. Green lines show the values obtained with standard regression analysis. The black line shows a slope of one.

Figure S.16: MRI, segmentations, simulations, recordings and fit for P016

# P017

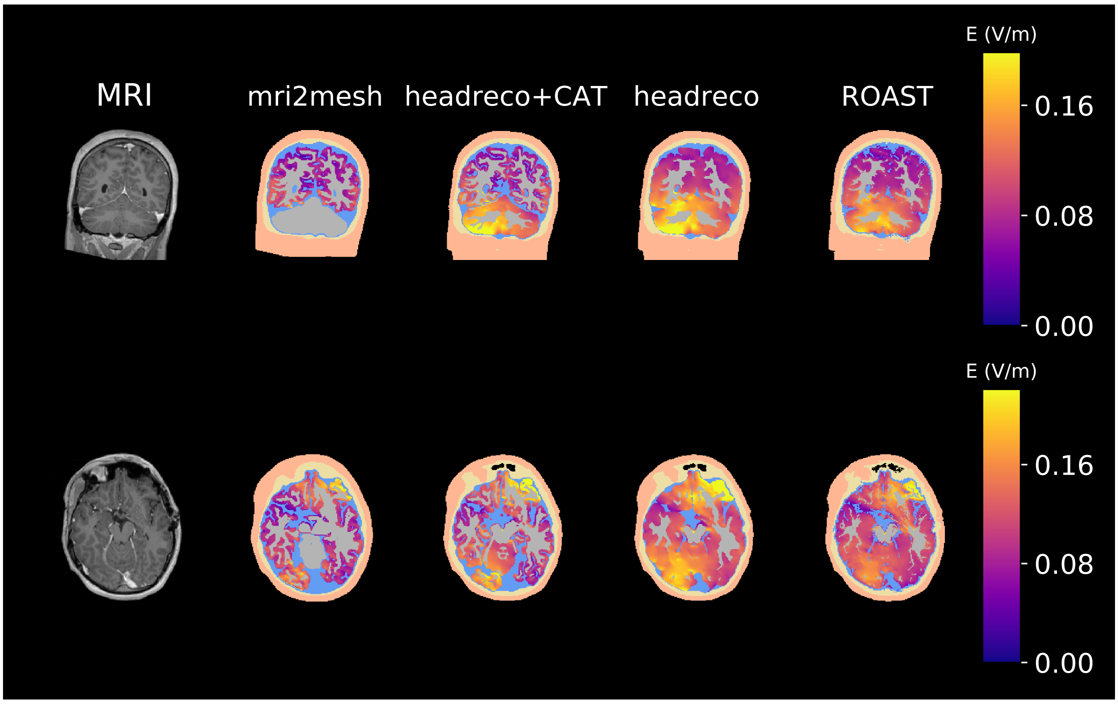


(a) T1-weighted image (first column), and segmentations with the electric field in gray matter

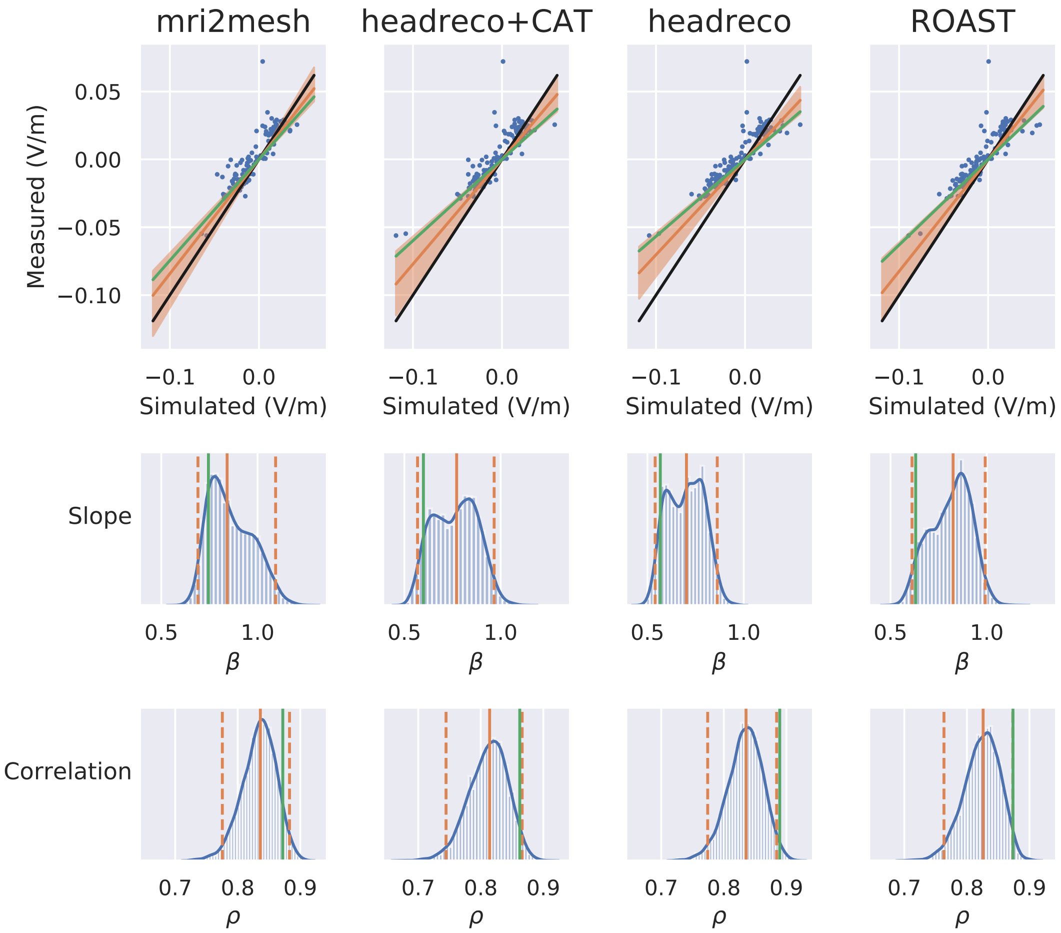


(b) Measured and simulated potential differences (first row), posterior probability for the slope $\beta$ (second row) and correlation $\rho$ (third row). Orange lines show the median and 95% compatibility interval obtained with the Bayesian errors-in-variables model. Green lines show the values obtained with standard regression analysis. The black line shows a slope of one.

Figure S.17: MRI, segmentations, simulations, recordings and fit for P017
